# A cellular and molecular analysis of SoxB-driven neurogenesis in a cnidarian

**DOI:** 10.1101/2022.03.22.485278

**Authors:** Eleni Chrysostomou, Hakima Flici, Sebastian G Gornik, Miguel Salinas-Saavedra, James M Gahan, Emma T McMahon, Kerry Thompson, Shirley Hanley, Michelle Kilcoyne, Christine E. Schnitzler, Paul Gonzalez, Andreas D Baxevanis, Uri Frank

## Abstract

Neurogenesis is the generation of neurons from stem cells, a process that is regulated by SoxB transcription factors (TFs) in many animals. Although the roles of these TFs are well understood in bilaterians, how their neural function evolved is unclear. Here, we use *Hydractinia symbiolongicarpus*, a member of the early-branching phylum Cnidaria, to provide insight into this question. Using a combination of mRNA *in situ* hybridization, transgenesis, gene knockdown, transcriptomics, and *in-vivo* imaging, we provide a comprehensive molecular and cellular analysis of neurogenesis during embryogenesis, homeostasis, and regeneration in this animal. We show that SoxB genes act sequentially. Stem cells expressing *Piwi1* and *Soxb1*, which have a broad developmental potential, become neural progenitors that express *Soxb2* before differentiating into mature neural cells. Knockdown of SoxB genes resulted in complex defects in embryonic neurogenesis. *Hydractinia* neural cells differentiate while migrating from the aboral to the oral end of the animal, but it is unclear whether migration per se or exposure to different microenvironments is the main driver of their fate determination. Our data constitute a rich resource for studies aiming at addressing this question, which is at the heart of understanding neurological disorders in all animals, including humans.

## INTRODUCTION

Neurogenesis − the generation of neuronal cells − is thought to be a partially conserved process in metazoans (Bosch et al., 2017; Gahan et al., 2017; Kelava et al., 2015; Rentzsch et al., 2016; Tournière et al., 2020; Watanabe et al., 2014) but the early evolutionary history of this process remains unclear. SoxB transcription factors (TFs) are central players in neurogenesis across a broad range of animal taxa including vertebrates, insects, nematodes, planarians, and cnidarians (Alqadah et al., 2015; Meulemans and Bronner-Fraser, 2007; Richards and Rentzsch, 2015; Ross et al., 2018) but how and when these genes became co-opted into the process of neuron formation is unknown. Vertebrate SoxB TFs are commonly subdivided into the SoxB1 group (Sox1, Sox2, and Sox3) that maintain stemness in neural stem cells, while SoxB2 group members (Sox14 and Sox21) are involved in neural differentiation (Bylund et al., 2003; Holmberg et al., 2008). Hence, vertebrate SoxB genes act sequentially in neurogenesis, but it is unclear whether this feature is ancestral or derived in the metazoan animals. Furthermore, SOX orthology has been notoriously hard to interpret and, therefore, a similar SoxB subdivision has not been observed in invertebrates (Flici et al., 2017; Jager et al., 2011; Richards and Rentzsch, 2015; Schnitzler et al., 2014). Finally, expression and functional data on SOX genes in the context of neurogenesis are still rare in non-bilaterians, making an evolutionary inference difficult to establish.

Cnidarians are the sister group to bilaterians (Kayal et al., 2018; Zapata et al., 2015) and, as such, they represent an interesting outgroup to study the evolution of neurogenesis. As in vertebrates, multiple SoxB genes are expressed in the developing nervous systems of representatives of the two main cnidarian clades, Anthozoa (corals and sea anemones) (Magie et al., 2005; Richards and Rentzsch, 2014, 2015) and Medusozoa (hydroids and jellies) (Flici et al., 2017; Jager et al., 2006). While we know that some cnidarian SoxB genes are required for neurogenesis, their function is not well understood. *Hydractinia symbiolongicarpus* is a colonial hydrozoan cnidarian (Figure S1) (Frank et al., 2020) and, like other hydrozoans, this animal possesses a population of stem cells, known as i-cells. These i-cells self-renew and contribute to somatic lineages, including neurons, and to germ cells throughout life (DuBuc et al., 2020; Gahan et al., 2016). The colonial nature of the animal enable i-cells to migrate throughout the colony and populate individual polyps. Hence, i-cells constitute a single population for the entire colony. i-cells first appear in embryonic endoderm (Gahan et al., 2016) but migrate to the outer tissue layer of the body, the epidermis, during metamorphosis (Figure S2), remaining there during adult life. In the mature animal, undifferentiated i-cells are found in interstitial spaces of the epidermis, primarily of the lower body column. They differentiate while migrating orally (Bradshaw et al., 2015), giving rise to various cell types including the two main neuronal lineages: neurons and nematocytes − the latter being specialized stinging cells that are used by cnidarians to capture prey or for defense from predators. The *Hydractinia* nervous system has a typical cnidarian nerve-net structure with a higher concentration of neurons at its oral end (Figures S3 & S4). The *Hydractinia* genome encodes three SoxB-like genes (*Soxb1, Soxb2,* and *Soxb3*) whose orthologous relation to specific bilaterian homologs cannot be resolved due to low sequence conservation. Hence, their names do not implicate them as being part of any bilaterian SoxB subgroup (Flici et al., 2017). A previous study has shown that *Hydractinia Soxb2* is expressed in proliferative, putative neural progenitors, and *Soxb3* is expressed in neural cells, identified by their distinctive morphology. Both genes are essential for adult neurogenesis and nematogenesis (Flici and Frank, 2018; Flici et al., 2017). However, the functions of *Soxb1*, *Soxb2*, and *Soxb3* in embryonic neurogenesis remain unknown.

Here, we have studied the function of all three *Hydractinia* SoxB genes in embryogenesis. To observe the dynamic expression of these genes in adults, we have generated *Soxb1* and *Soxb2* transgenic reporter animals and tracked individual cells during differentiation using *in vivo* time-lapse imaging. We also used fluorescence-activated cell sorting (FACS) and droplet microfluidics approaches (InDrop-seq) as a first step towards generating cell type-specific and single-cell transcriptomes, then analyzed the cell cycle characteristics of *Hydractinia* cell fractions. We find that SoxB genes act sequentially in adults as i-cells differentiate into neurons and nematocytes. Knockdown experiments indicate a complex role for SoxB genes in the differentiation of embryonic neuronal cells. Under *Soxb2* knockdown conditions, we have documented biased differentiation to specific neural cell types. Our study provides insight into the evolution of SoxB genes and their function in neurogenesis across Metazoa.

## RESULTS

### Expression pattern of SoxB genes in *Hydractinia*

We studied the expression pattern of all three SoxB genes in *Hydractinia* using both traditional, tyramide amplification-based fluorescence mRNA *in situ* hybridization (FISH) and Signal Amplification By Exchange Reaction (SABER) single molecule FISH (Kishi et al., 2019). In sexual polyps, *Soxb1* was expressed in cells morphologically resembling i-cells and germ cells in males and females (Figure 1A-D). In feeding polyps, *Soxb1* was expressed in cells that resembled i-cells both with respect to their morphology and their anatomical location, being primarily distributed in the lower body column of the polyp and largely absent from the oral end of the animal (Bradshaw et al., 2015) (Figure 1E-G). To confirm the identity of *Soxb1*-expressing cells, we used double FISH with *Piwi1*, a known marker for *Hydractinia* i-cells and germ cells (DuBuc et al., 2020; Gahan et al., 2016). We found that all *Soxb1*^+^ cells also expressed *Piwi1* and vice versa (Figure 1). Therefore, we conclude that *Soxb1* is expressed in i-cells, i.e. in stem cells that give rise to somatic cells (including neurons and nematocytes) and to germ cells. *Soxb2* expression partially overlapped with *Soxb1* in the mid-region of the polyp body column, being largely absent from the lower (aboral) area of the polyp that primarily included only *Piwi1^+^/Soxb1*^+^ cells (Figure 2A-C). *Soxb3* was expressed in the middle and upper (i.e., oral) parts of the polyp and partly overlapped with *Soxb2* in the mid-region (Figure 2D-F). These findings are consistent with a previous study that investigated *Soxb2/Soxb3* expression (Flici et al., 2017). It is also in line with the known pattern of distribution of *Hydractinia* cells, with i-cells being restricted to the lower body column, and neural cells mainly concentrated in the oral pole.

**Figure 1.**
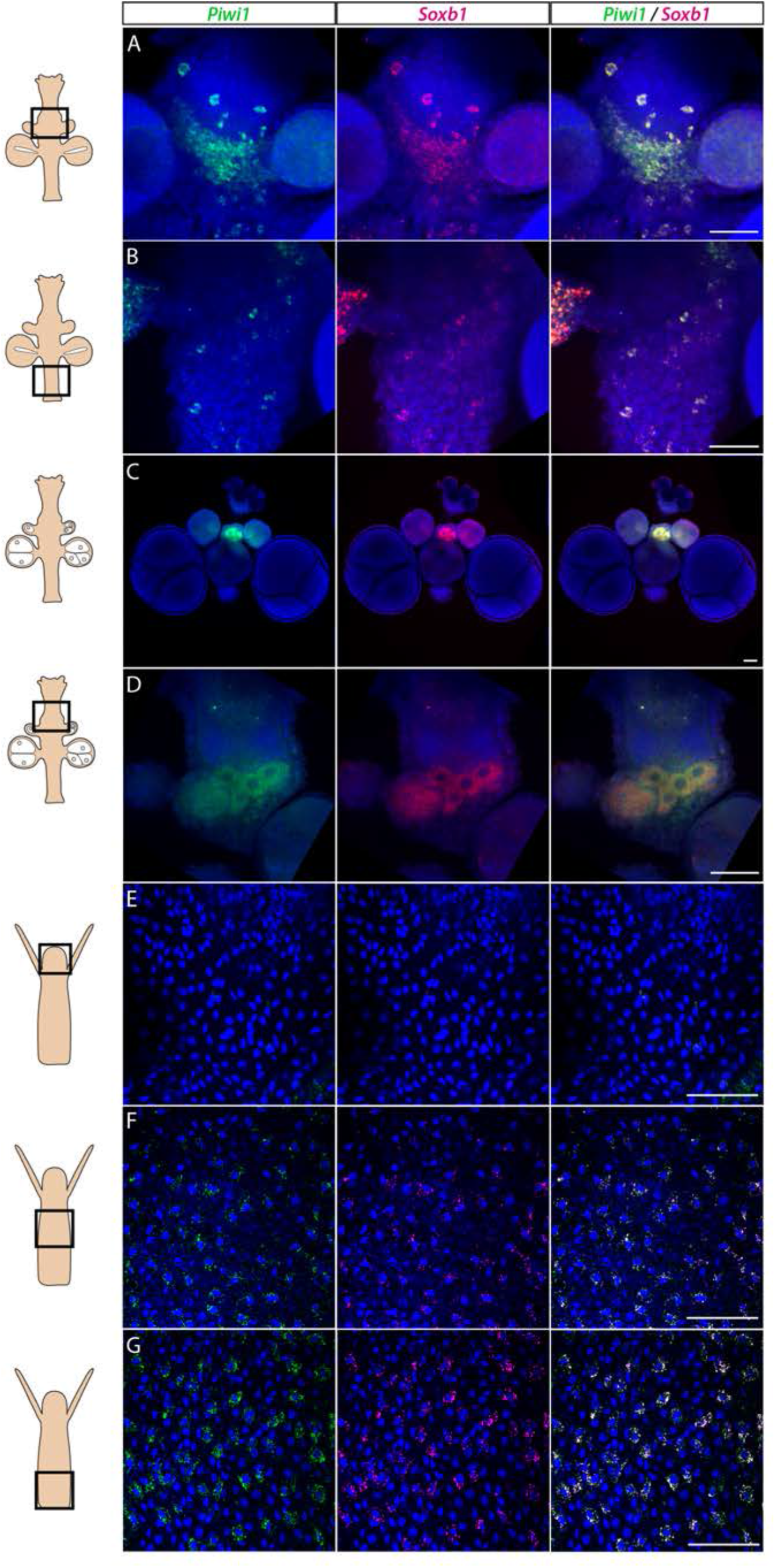
*Piwi1* and *Soxb1* are co-expressed in i-cells and germ cells. (A-D) Analysis of sexual polyps’ mid (A) and lower (B) body column, and early (C) and late (D) oocytes. (E-G) Analysis of feeding polyps head (E), mid-(F) and lower (G) body column. (A-D) were tyramide-based FISH; (E-G) were single-molecule FISH. Scale bars 40 μm.

**Figure 2.**
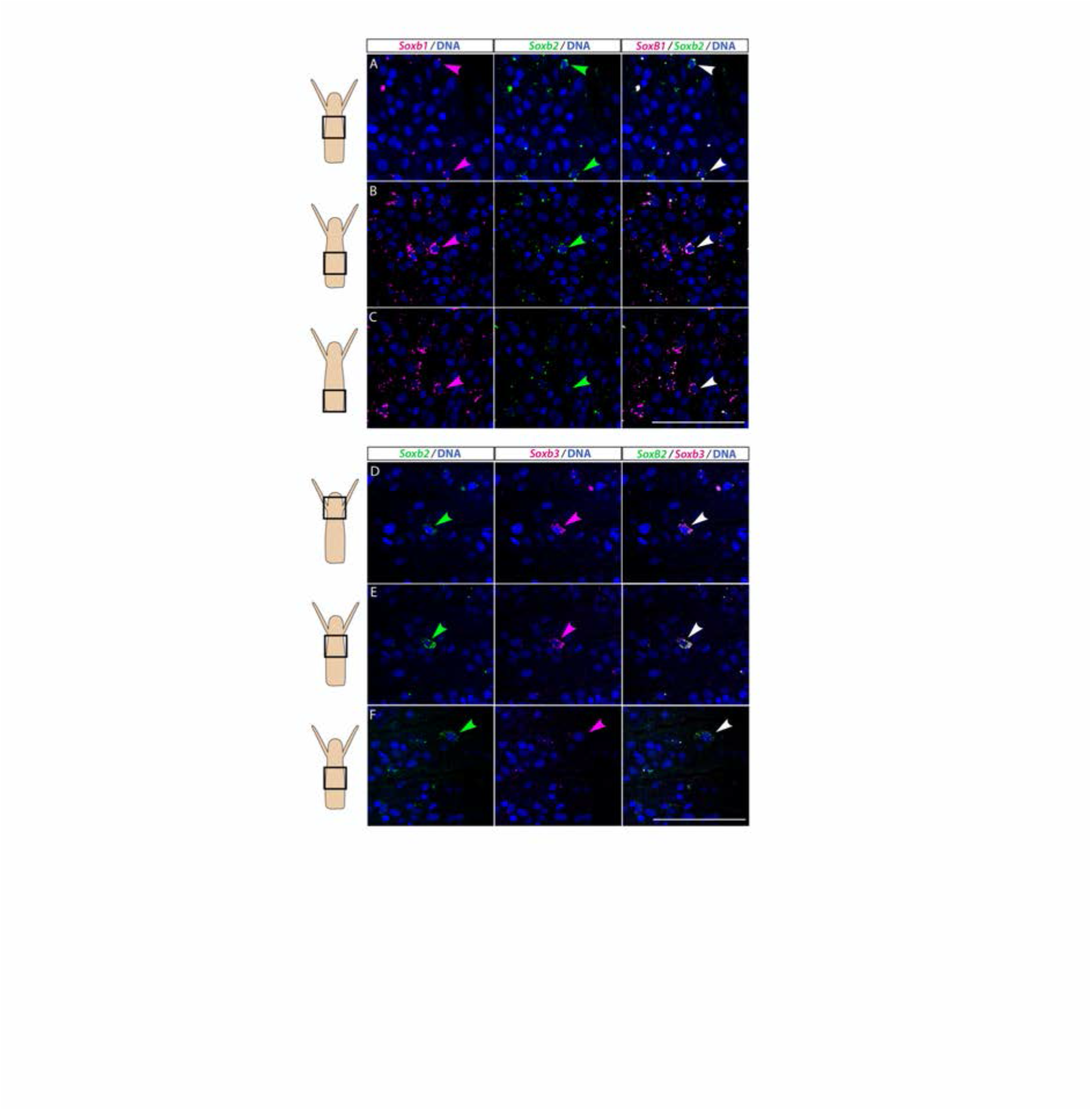
Partial overlap in the expression of *Soxb1* and *Soxb2*, and *Soxb2* and *Soxb3* in feeding polyps. (A-C) Single molecule FISH with probes against *Soxb1* and *Soxb2 showing* upper-mid (A), lower-mid (B), and lower (C) body column. (D-F) Single molecule FISH with probes against *Soxb2* and *Soxb3* showing the head region (D), upper-mid (E), and lower-mid (F) body column. Arrowheads point to double-positive cells. Scale bars 40 μm.

While mRNA FISH experiments, described above, informed us about cells that expressed SoxB genes in space and time at cellular resolution, they did not provide direct information on the fate of these cells. To obtain these data, we generated transgenic reporter animals that expressed tdTomato or GFP, driven by the endogenous *Soxb1* or *Soxb2* genomic control elements, respectively (Figure 3A-I). Attempts to generate a *Soxb3* reporter were unsuccessful. Instead, we used an *Rfamide* reporter animal that expresses GFP in RFamide^+^ neurons (Figure 3J-L). The rationale of this approach was that the long half-life of fluorescent proteins would enable tracking cells even after their fate is changed and the reporter gene shut down. Fluorescence of the *Soxb1::tdTomato* reporter was dim *in vivo* but the signal was enhanced in fixed animals through the use of anti-RFP antibodies (Figure 3A-E). In contrast, the signal produced by the *Soxb2* and *Rfamide* reporters, both expressing GFP, was sufficiently bright to be viewed *in vivo* using fluorescence stereomicroscopy (Figure 3F-L).

**Figure 3.**
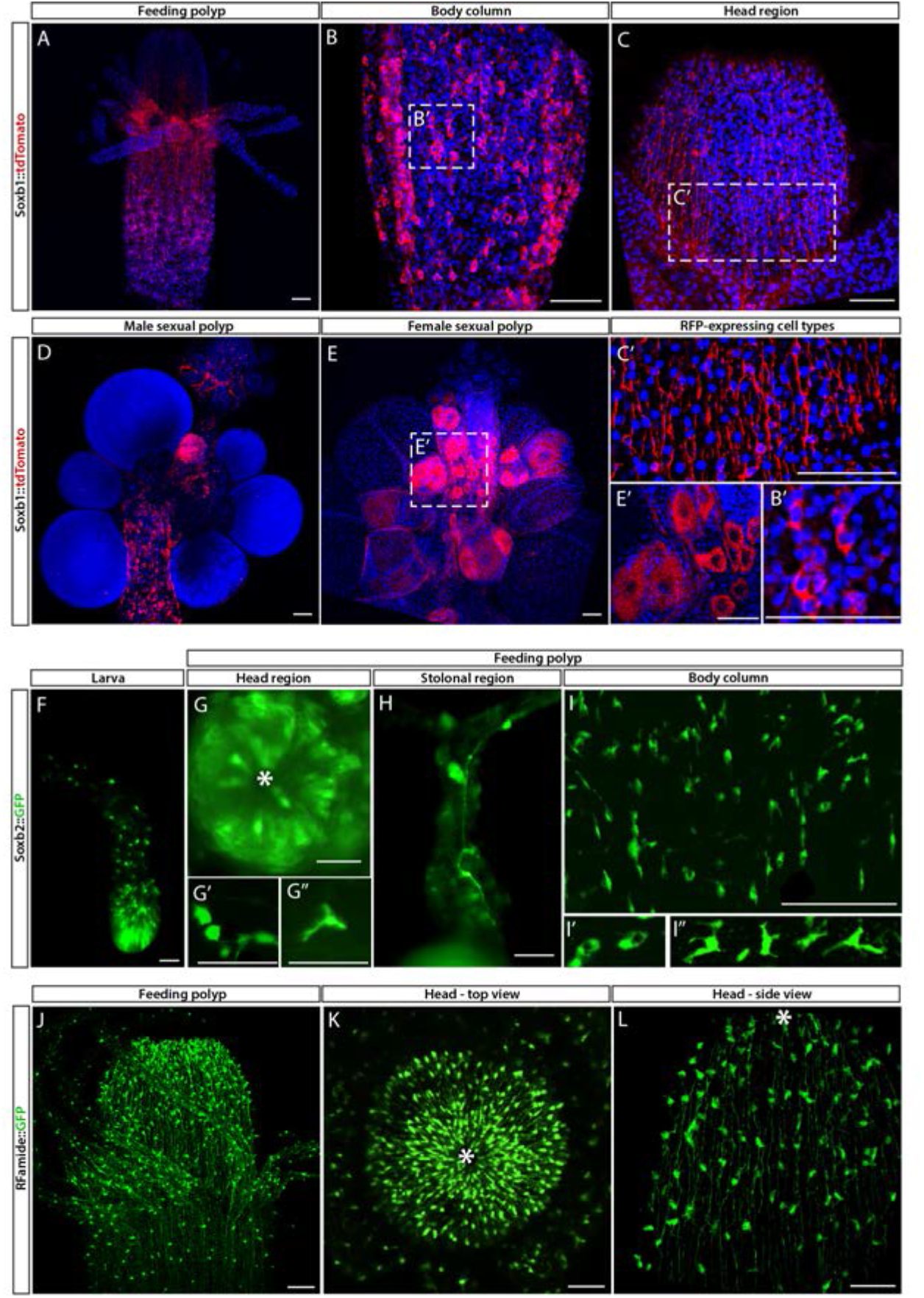
*Soxb1, Soxb2,* and *Rfamide* transgenic reporter animals. (A-E) *Soxb1::tdTomato* reporter animal. Animals were fixed and stained with an anti-dsRed antibody. (A) Whole feeding polyp. (B) Lower polyp body column. (B’) higher magnification of tdTomato^+^ i-cells. (C) Head region including base of tentacles. (C’) close-up showing tdTomato^+^ neurites. (D) Male sexual polyp. (E) Female sexual polyp. (E’) Higher magnification of tdTomato^+^ oocytes. (F-I) Live imaging of a *Soxb2::GFP* reporter animal. (F) planula larva. (G) Oral pole of a polyp viewed from above. (G’, G’’) Higher magnification of GFP^+^ cells with neural morphology. (H) GFP^+^ interconnected neurons. (I) Mid-body column region of polyp showing GFP^+^ cells. (I’) GFP^+^ nematoblasts (based on the presence of a nematocyst capsule). (I’’) GFP^+^ neurons. (J-L) Live imaging of *Rfamide::GFP*^+^ reporter feeding polyps. (J) Upper polyp region including head and tentacles. (K) Oral pole of feeding polyp viewed from above. (L) Higher magnification of the head region, showing GFP^+^ neural network. Asterisk point to oral ends. Scale bars 40 μm.

Contrary to the mRNA FISH data (Figure 1), the *Soxb1* promoter-driven tdTomato was not only observed in stem/progenitor cells but was present in neural cells as well, based on the distinct morphology of these cells (Figure 3A-C). In sexual polyps, tdTomato was also observed in maturing germ cells (Figure 3D and E). This shows that *Soxb1*^+^ cells differentiated into neural cells and gametes as expected. The *Soxb2* promoter-driven GFP was visible in neurons and nematocytes but not in i-cells (Figures 2 and 3F-I), suggesting that *Soxb2*^+^ cells were more committed than i-cells (i.e. *Soxb1*^+^ cells). The spatial distribution of *Soxb1^+^* and *Soxb2*^+^ cells was distinct from that of their respective progeny, consistent with migrating neural progenitors that differentiate during and following their migration away from the lower body column area where i-cells and their early progeny reside. The *Rfamide* reporter was only visible in sensory and ganglionic neurons (Figure 3J-L), consistent with the expression pattern of this gene in neurons but not in neural progenitors (Gahan et al., 2017; Kanska and Frank, 2013).

FISH expression patterns (Figure 2) showed partial overlap between *Soxb1* and *Soxb2*, as well as between *Soxb2* and *Soxb3*, the latter confirming previous reported observations (Flici et al., 2017). Furthermore, analysis of transgenic reporter animals indicated that *Soxb*1^+^ and *Soxb2*^+^ cells give rise to neural cells (Figures 2 and 3). To confirm within-lineage transitions between *Soxb1* and *Soxb2* expression, we generated double *Soxb1*:*:tdTomato* / *Soxb2::GFP* reporter animals by crossing a *Soxb1*:*:tdTomato* female with a *Soxb2::GFP* male. The *Soxb1*- driven tdTomato fluorescence that was too dim to be viewed live by stereomicroscopy, was readily visible *in vivo* using a more sensitive spinning disk confocal microscope. Double transgenic animals were mounted in low-melt agarose and subjected to prolonged *in vivo* spinning disk confocal imaging. We observed tdTomato-positive (i.e., *Soxb1*^+^), GFP-low or negative (i.e., *Soxb2*^-^) cells transforming to double positive cells (Figure 4; Figure S5). Collectively, our mRNA expression data and the ones obtained previously (Flici et al., 2017), together with the results of our live imaging of transgenic reporter animals are consistent with sequential expression of SoxB genes in the neural lineage.

**Figure 4.**
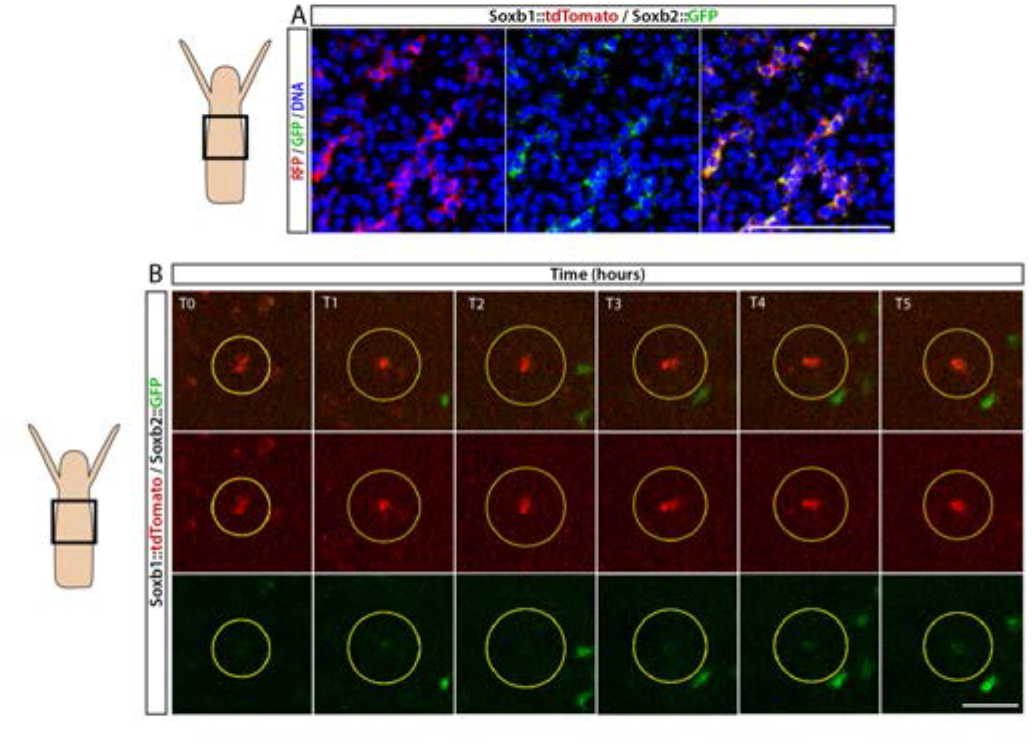
Sequential expression of *Soxb1* and *Soxb2*. (A) Immunofluorescence of *Soxb1::tdTomato* and *Soxb2::GFP* double positive cells that resemble differentiating neurons by morphology. (B) In vivo time lapse imaging of a *Soxb1::tdTomato*^+^ cells (shown in red) with increasing levels of *Soxb2::GFP* (shown in green) over a timeframe of 8 hours (T0=3 hpd; T1=4 hpd… T5=8 hpd). Green *Soxb2::GFP* cells that appear in the image probably migrated into the confocal plane during imaging. Scale bars 40 μm.

### Cellular and molecular characterization of *Hydractinia* neural cells

We used a recently developed flow cytometry/FACS protocol (DuBuc et al., 2020) to analyze and physically sort cells from dissociated transgenic reporter animals. The long half-life of the fluorescent reporter proteins resulted in the isolation of not only cells that expressed the reporter genes but also their progeny. To minimize the contribution of progeny to the sorted cellular fraction, we used a FACS gating strategy to select live, nucleated single cells and sorted populations that had low side scatter characteristics and high levels of GFP (Figure 5A-C and S6).

**Figure 5.**
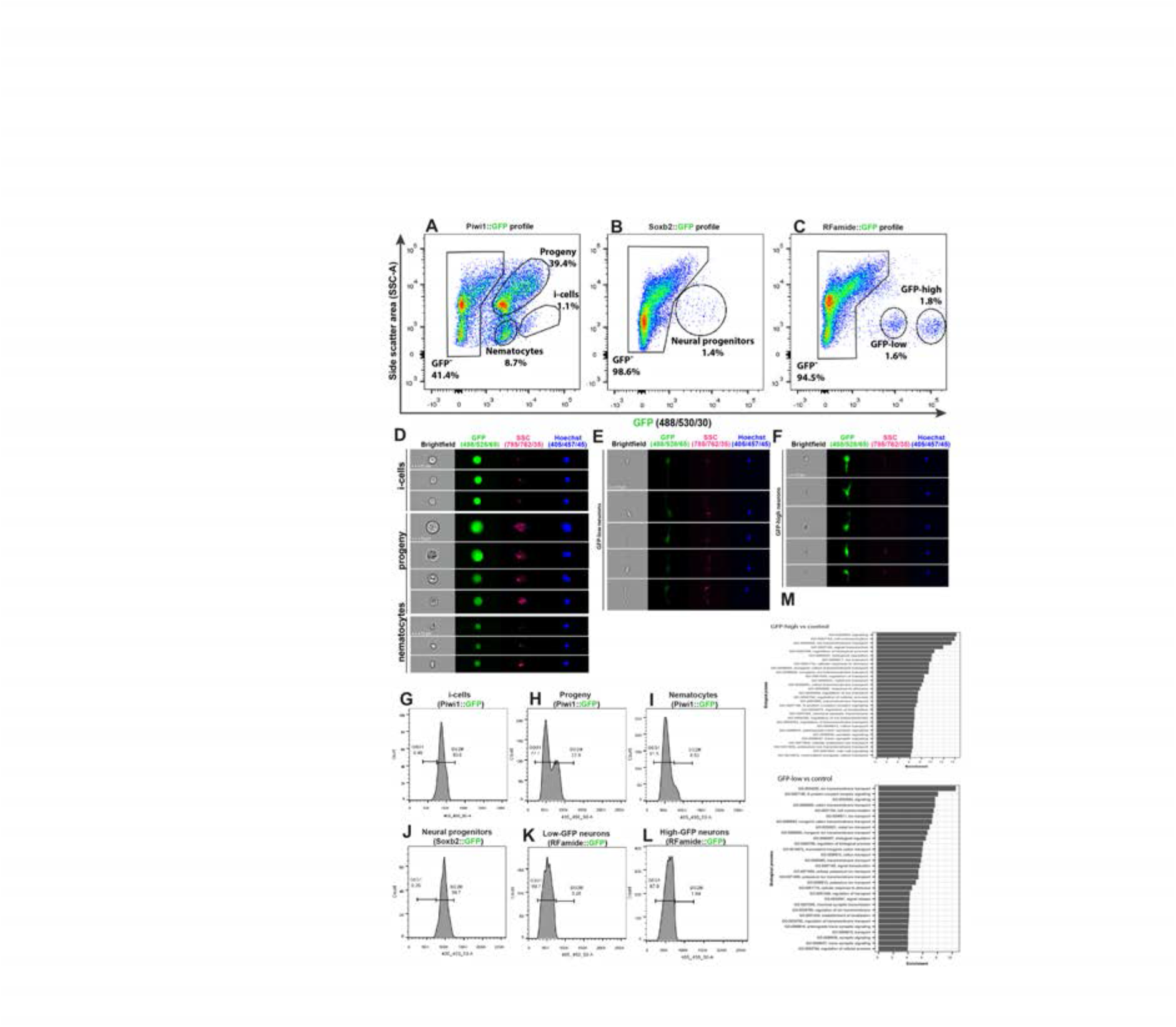
Analysis of dissociated *Hydractinia* cells. (A) Gating strategy of i-cells, nematocytes, mixed progenitors, and GFP^-^ cells from a *Piwi1* reporter animal. (B) Gating of putative neural progenitors from a *Soxb2* reporter animal. (C) Gating two distinct populations of *Rfamide*::GFP+ neurons. (D-F) Imaging flow cytometric analysis of cells from the transgenic reporter animals. (G-L) Cell cycle analysis of distinct, sorted cell populations. (G) i-cells are mostly in S/G2/M. (H) i-cell progeny are distributed along the cell cycle. (I) Nematocytes are mostly in G1. (J) *SoxB^+^* cells (putative neural progenitors) are mostly in S/G2/M. (K) Low and (L) high *Rfamide*^+^ neurons. (M) Differential gene expression analysis of sorted cell populations. Genes expressed by *Rfamide*^+^ neurons and *Soxb2*^+^ cells are listed.

A previously established *Piwi1*::*GFP* reporter animal (Bradshaw et al., 2015) was used to selectively isolate i-cells by FACS rather than the *Soxb1::tdTomato* reporter animal. tdTomato fluorescent protein is optimally excited by a 561nm yellow/green laser; the configuration of the BD FACS Aria cell sorter in the NUI Galway flow cytometry core facility does not include a 561nm laser. The usage of *Piwi1*::*GFP* to isolate i-cells was possible given that the two genes have been shown to be co-expressed in i-cells (Figure 1). *Piwi1^+^* (i.e., *Soxb1*^+^ i-cells) are known to represent a rare population, representing approximately 1.4% of the live single cell population of the *Soxb2::GFP* transgenic animal (DuBuc et al., 2020). We found that *Soxb2*^+^ cells (i.e., putative neural progenitors) were an equally rare population (Figure 5A-B). *Rfamide*^+^ neurons were distributed among two distinct populations that differed in levels of GFP expression, accounting for 1.6% and 1.8% of the live single cells, respectively (Figure 5C). Imaging flow cytometry of the *Piwi1*::GFP reporter animal (i.e., cells that also express *Soxb1*) revealed that the brightest cells were small with a high nuclear to cytoplasmic ratio, consistent with stem cell morphology (Figure 5D). The two Rfamide^+^ cell populations did not differ morphologically based on imaging flow cytometry except in their GFP expression (Figure 5E, F); their exact nature is yet unclear. Flow cytometric cell cycle analysis (Figure 5G-L) revealed that the *Piwi1^+^* i-cells were almost exclusively in S/G2/M phase (Figure 5G), similar to what has been observed in i-cells from the freshwater polyp *Hydra* (Buzgariu et al., 2014). However, i-cell progeny, defined by low GFP fluorescence and constituting a highly heterogeneous population, were found in all stages of the cell cycle (Figure 5H). *Soxb2*^+^ cells that include putative neural progenitors (Flici et al., 2017), had a cell cycle profile similar to i-cells, mostly found during the S/G2/M phases of the cell cycle (Figure 5J). *Rfamide*^high^ and *Rfamide*^low^ neurons that are thought to be terminally differentiated, were mostly in G0/G1, similar to what was observed in nematocytes (Figure 5I, K-L). The robustness of the flow cytometric analysis was demonstrated by studying other cellular fractions including epithelial cells from an epithelial reporter (Künzel et al., 2010), male germ cells (from the *Piwi1* reporter), and cells from an *Ef1a* knock-in animal where GFP is expressed ubiquitously (Sanders et al., 2018) (Figures S6-S7).

We used FACS to physically isolate the following cell fractions: *Soxb2*^high^, *Rfamide*^high^, and *Rfamide*^low^ neurons. We generated transcriptomes for each of these cell fractions and used a previously published i-cell transcriptome (DuBuc et al., 2020). We then compared gene expression between the combinations shown in Table S.1. Given the long half-life of fluorescent proteins, the *Soxb2*^+^ cells transcriptome we present (Supplemental File 1) probably includes also early stages of their differentiation. The two RFamide^+^ neuronal populations are probably different, as the RFamide neuropeptide is considered a marker for differentiated neurons. Indeed, typical neuronal genes were expressed by these cells (Figure 5M; Supplemental File 1). The RFamide^+^ neurons fraction also expressed minicollagen genes that are markers of nematogenesis. Co-expression of minicollagens and precursors of neuropeptides has been reported previously (Sunagar et al., 2018; Wolenski et al., 2013) so this observation in RFamide^+^ cells is not surprising given that nematocytes are neural cells.

To provide further insight into the composition of the neural cells population, we used a droplet microfluidic single cell handling system (InDrop) (Klein et al., 2015) to generate a single-cell transcriptome dataset based on 7,071 cells from *Hydractinia* wild type feeding polyps. This allowed us to identify 22 cell clusters (Figure S8A) and their associated upregulated marker genes (Figure S8B); this may be an underestimation of the cellular complexity of *Hydractinia* as compared to that of other cnidarians (Sebe-Pedros et al., 2018; Siebert et al., 2019). However, we were able to identify putative i-cells, nematoblasts, gland cells, neurons, and a group of mixed progenitors. Upregulated marker gene expressions generally recapitulated the expression profiles of the physically sorted cells from the transgenic reporter animals, with up to 15% of markers corresponding to DEG transcript of the sorted cell types (Figure S8C; Supplementary File 2). Specifically analyzing the expression of *Soxb1-3* (Figure S8D), we find that both *Soxb1* and *Soxb2* are expressed in a subset of i-cells; these cells are presumably in transition to becoming *Soxb2*-only positive progenitor cells (clusters 13 and 14 in Figure S8A). Finally, *Soxb3* is only expressed in few cells in cluster 18 (Figure S8A and S8D). It also appears that subpopulations of neural cells transition from a *Soxb2* RFamide low state (cluster 14) that express a G-protein coupled receptor for glutamate (Figure S8E) to a *Soxb2* RFamide high state (cluster 13), of which some eventually become *Soxb2* negative while still expressing high levels of RFamide (cluster 11). These are presumably terminally differentiated neurons as determined by the marker genes Kaptin and Contactin2 (Figure S8E). This is consistent with the downregulation of *Soxb2* resulting in near absence of RFamide^+^ neurons (Figure 7) and bolsters the role of *Soxb2* as a marker for neural progenitor cells. Combined, the bulk- and-single cell analyses provide a valuable resource for further analyses of cnidarian cells, allowing for the extended analysis of nematoblasts as a way to characterize both known and novel marker genes (Figure S8F).

### *Hydractinia* head regeneration involves *de novo* neurogenesis

*Hydractinia* can regenerate a decapitated head that includes a fully functional nervous system within 2-3 days (Bradshaw et al., 2015). This predictable generation of many new neurons provides an opportunity to study neurogenesis in regeneration. To identify the cellular source of regenerative neurogenesis, we first characterized the dynamics of nervous system regeneration over 72 hours using *Rfamide::GFP* reporter transgenic animals (Figure 6). Animals were decapitated and allowed to regenerate their heads. Subsets of these animals were then fixed at 12, 24, 36, and 48 hours post-decapitation (hpd). Prior to fixation, they were incubated for 30 minutes in EdU. Fixed samples were stained for EdU, GFP (reporting *Rfamide*), and Piwi2 (an i-cell marker; for anti-Piwi2 antibody validation; see Figure S9). We also analyzed intact animals that showed a well-developed head nervous system but had neither S-phase cells nor i-cells in their heads (Figure 6A), as expected (Bradshaw et al., 2015). At 12 hpd, there were many cycling cells in the stump area, few neurons (likely remnants of the amputated tissue), and few migrating Piwi2^low^ cells that are probably i-cell progeny (Figure 6B). At 24 hpd, a clear proliferative blastema had been established at the oral most pole, with i-cells (Piwi2^high^) and their progeny being numerous in the area. However, only a few RFamide^+^ neurons were visible (Figure 6C). At 36 hpd, new neurons were visible in the regenerating head and proliferative i-cells and progeny were still present (Figure 6D). Finally, at 48 hpd, a nearly complete head nervous system had been established and the regenerating head contained lower numbers of i-cells and progeny (Figure 6E). Hence, Piwi2^low^ cells, which are presumably i-cell progeny, were the first to migrate to the injury area, followed later on by Piwi2^high^ cells, which are probably stem cells. This is similar to what has been observed in planarian head regeneration, where neoblast progeny arrive in the injury area before neoblasts (Abnave et al., 2017). Of note, none of the existing neurons incorporated EdU during regeneration, consistent with neurons being terminally differentiated. To study nervous system regeneration *in vivo*, we used the *Rfamide*::GFP transgenic reporter animals to track individual neurons over the course of regeneration. The animals were decapitated, mounted in low-melt agarose, and imaged every hour for either the first 24 hours, or from 24-72 hpd. In all cases, the neurons remained stationary and did not proliferate during the observation period while a new head nervous system regenerated (Figure S10). Hence, similar to *Hydra* (Miljkovic-Licina et al., 2007), head nervous system regeneration in *Hydractinia* involves *de novo* neurogenesis rather than proliferation or migration of existing neurons.

**Figure 6.**
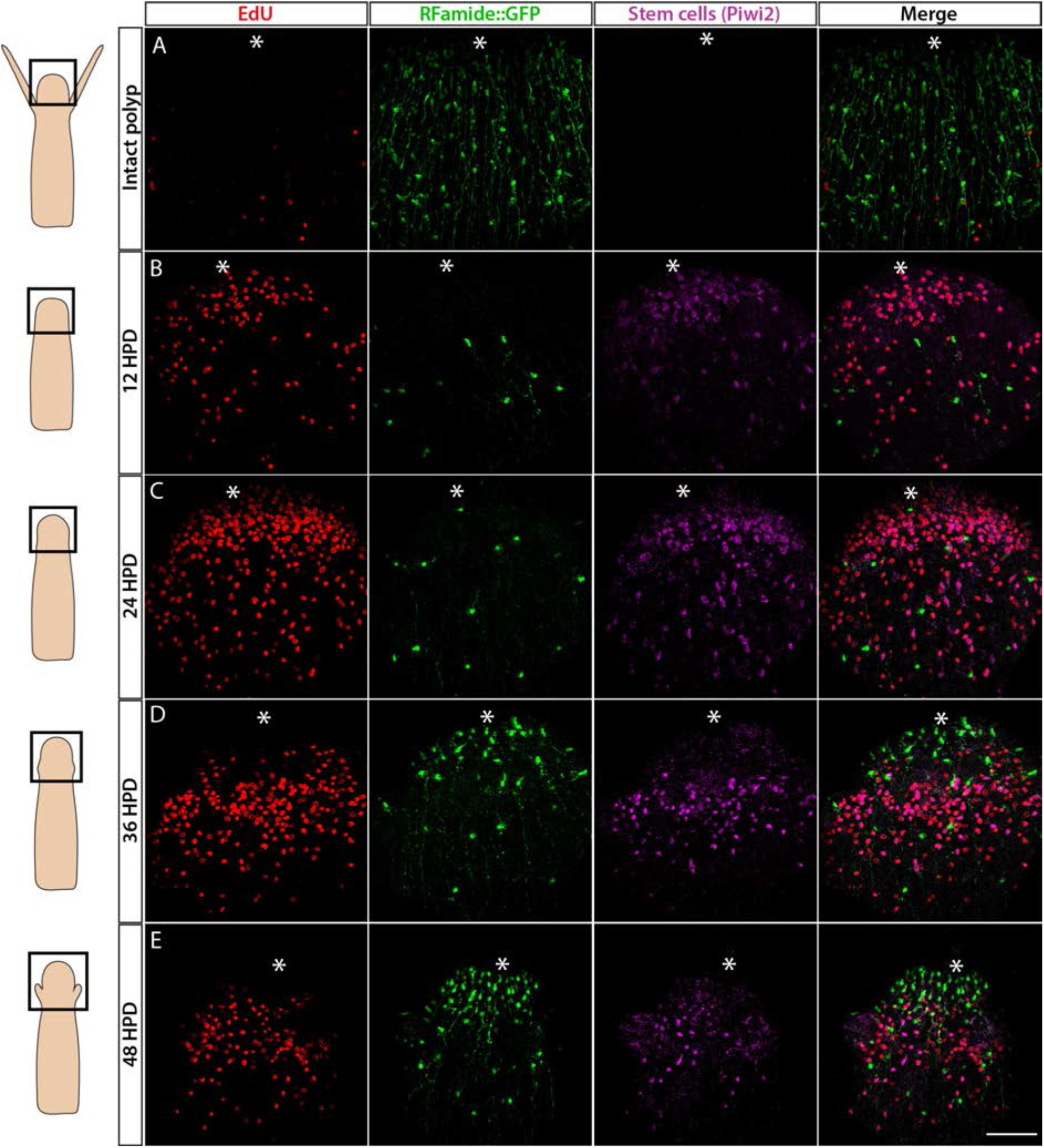
Cellular events during *Hydractinia* head regeneration in an Rfamide::GFP transgenic reporter animal. (A) Intact head, characterized by few S-phase cells, extensive RFamide^+^ neuronal network, and no Piwi1^high^ i-cells or their putative Piwi1^low^ progeny. (B) 12 hours post decapitation (hpd) showing many proliferative cells, only few remnant RFamide ^+^ neurons, and few immigrating i-cells and progeny. (C) 24 hpd. High numbers of EdU incorporating cells, few RFamide^+^ neurons, and increasing i-cells and putative progeny are seen at the oral regenerating end. (D) 36 hpd. High number of S-phase cells, increased number of RFamide^+^ neurons, and further increase in i-cell numbers are seen. (E) 48 HPD Decreasing numbers of S-phase cells, increasing numbers of RFamide^+^ neurons and decreasing numbers of i-cells are observed. Scale bars 40 μm.

### SoxB genes are required for embryonic neurogenesis

It has been shown previously that *Soxb2* and *Soxb3* are essential for neurogenesis in adult *Hydractinia* (Flici et al., 2017). However, the role of *Soxb1* has not been studied, nor has the role of these TFs in other life stages. Hence, we set out to study the role of all three SoxB genes in embryogenesis. For this, we used short hairpin RNA (shRNA)-mediated knockdown by microinjection into zygotes as previously described (DuBuc et al., 2020). The specificity of the treatment was assessed by FISH (Figure S11A). The animals were fixed at 72 hpf; at this stage of development, *Hydractinia* normally reaches the metamorphosis-competent planula larva stage.

*Soxb1* knockdown affected many aspects of larval development. First, the animals were on average smaller and underdeveloped (Figure S11B). Second, these animals had fewer Piwi1^+^ i-cells and numbers of neurons and nematoblasts was lower than normal. RFamide neuron numbers were low or absent altogether (Figure 7). Third, these larvae had defects in ciliation (Figures 7 & S12) and, as a result, their movement was restricted (larvae swim by coordinated cilia beat). Finally, the larvae had lost their ability to metamorphose upon CsCl treatment (Figure 7), with the latter probably being the result of a loss of neurons expressing GLWamide, the internal regulator of metamorphosis in *Hydractinia* (Gajewski et al., 1996; Müller and Leitz, 2002; Schmich et al., 1998). However, the number of S-phase cells was upregulated compared with the control and proliferative cells were also detectable in the epidermis; normally, these cells are restricted to the gastrodermis in larvae (Figure 7). The above defects could be explained by *Soxb1* playing a role in maintaining stemness in i-cells, similar to the role of Sox2 in mammalian pluripotency (Boyer et al., 2005; Takahashi and Yamanaka, 2006). The loss of i-cells, as observed by Piwi1 staining (Figure 7), would be expected to directly affect all i-cell derivatives that include neurons and nematocytes, as well as other cell types. The increased number of S-phase cells could reflect a compensatory response to loss of i-cells.

**Figure 7.**
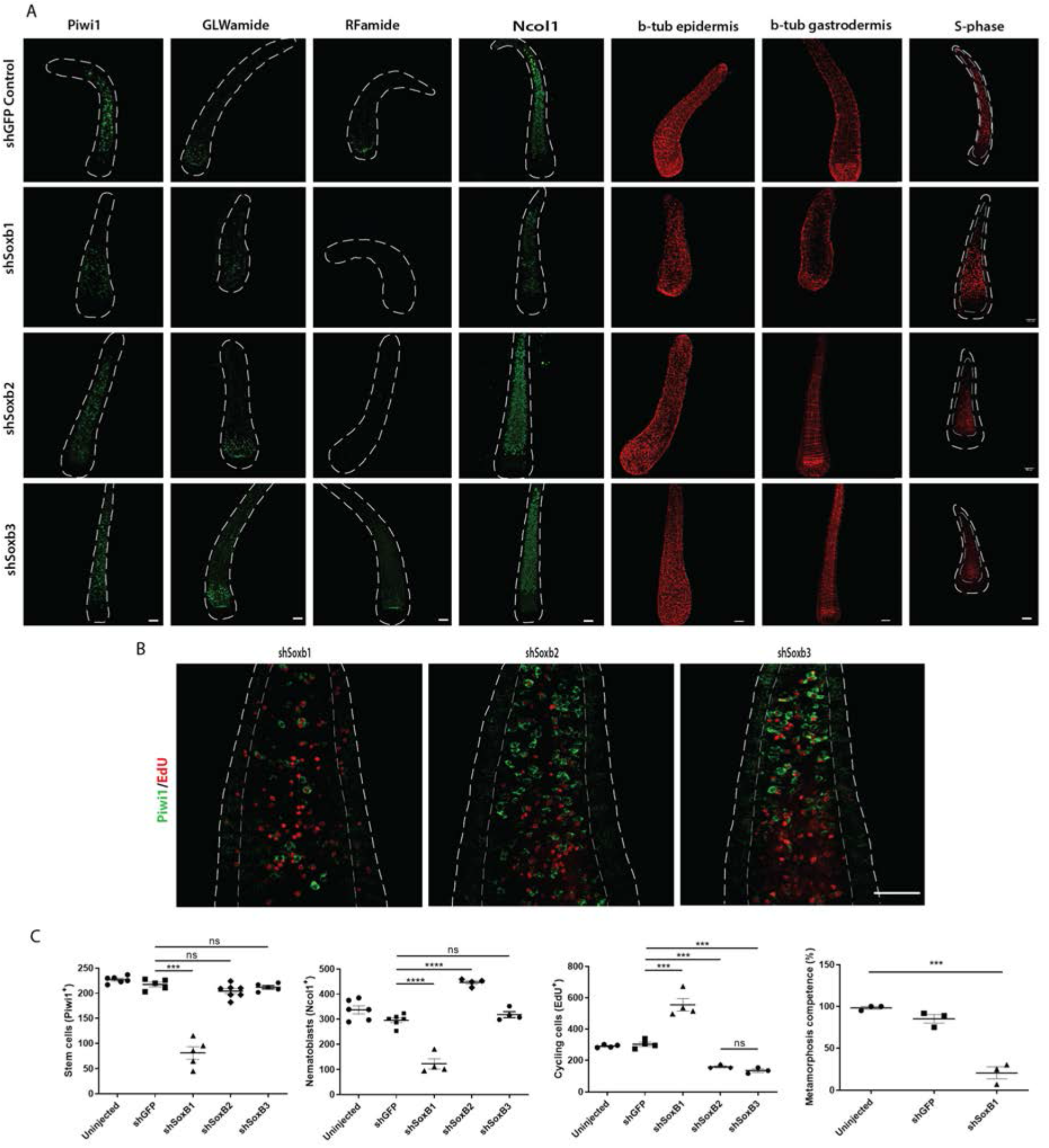
Effects of SoxB genes downregulation in embryogenesis. (A) shRNA-mediated knockdown of *Soxb1*, *Soxb2*, and *Soxb3*. (B) Higher magnification of Piwi1^+^ and S-phase cells showing reduced i-cell numbers following *Soxb1* downregulation. Note proliferative cells in epidermis, not seen in untreated and following *Soxb2* or *Soxb3* downregulation. (C) Quantification of phenotypes.

Downregulation of *Soxb2* had no visible effect on i-cells or on GLWamide^+^ neurons (Figure 7) but RFamide^+^ neurons were nearly absent. Nematoblast numbers had increased and no significant difference was observed on cycling cells. These phenotypes are markedly distinct from what is seen after downregulation of *Soxb2* in adult neurogenesis, where the numbers of neurons, nematocytes, and cycling cells are noticeably reduced (Flici et al., 2017). This could be explained by preferential differentiation of neural progenitors to GLWamide^+^ neurons over RFamide^+^ neurons if progenitors are limiting, given the essential role of the latter in metamorphosis. Alternatively, the SoxB2 protein might be maternally loaded or dispensable for embryonic GLWamide^+^ neurons. The increase in nematoblast numbers following *Soxb2* downregulation could indicate a nematogenesis-inhibiting role for this gene. Surprisingly, downregulation of *Soxb3* had no major effect on the animals, in contrast to the role of this gene in adult neurogenesis (Flici et al., 2017). The above data suggest an essential and complex role of *Soxb2* for embryonic neurogenesis, but also highlight major differences between embryonic and adult neurogenesis.

## DISCUSSION

Multiple SoxB proteins act to generate neurons across metazoans but orthologous relationships between individual SoxB members in different animal clades have been difficult to establish (Flici et al., 2017). This is either because multiple duplication events of an ancestral SoxB gene occurred within lineages, resulting in paralogous genes in any one clade, or that sequence drift over evolutionary time has blurred discernable orthologous relationships. However, the requirement for more than one SoxB gene to generate neurons in animals as distantly related as *Hydractinia*, *Drosophila*, and mice suggests that more than one SoxB gene had already been present before the cnidarian-bilaterian split to generate neurons in their last common ancestor; it does not exclude further lineage specific diversification and gene losses thereafter. Finally, it indicates a highly conserved mechanism of subfunctionization following gene duplication at the base of animals.

*Soxb1* is an i-cell marker and probably fulfills a role in stemness. Its downregulation resulted in loss of i-cells and impacted other aspects of development including the ability to metamorphose, which was expected given that i-cells provide progenitors to multiple lineages. This function of *Soxb1* is similar to the role of vertebrate Sox2. However, vertebrate Sox2 also acts in neural stem cells while in *Hydractinia*, there is no direct evidence so far to support a similar function. Furthermore, a population of long-term, self-renewing neural progenitors has yet to be identified in cnidarians. Our results from *Hydractinia*, along with data obtained from other cnidarians (reviewed by (Rentzsch et al., 2016), are also consistent with continuous generation of neurons from multi- or pluripotent cells that also contribute to non-neural lineages, without the existence of a long-term, self-renewing neural stem cell.

*Soxb2* was expressed in a population of cells with uncommitted morphology that differentiated to neural cells, based on GFP retention in neurons and nematocytes in the *Soxb2* reporter animal. Our *in vivo* analysis of *Soxb1/Soxb2* double transgenic reporter animals has shown that *Soxb1*^+^ cells transform into *Soxb2*^+^ cells (Figures 4 and S5). Downregulation of *Soxb2* resulted in neural-specific defects in embryos, consistent with this gene being expressed preferentially in the neural lineage (Figure 3). The latter is also supported by the neural transcriptional fingerprint (Figure 5M; Supplementary File 1) and single cell transcriptomic profiles of *Soxb2*^+^ cells (Figure S8C-S8E). Of note, deviation from normal development following *Soxb2* perturbation was not a simple reduction in all neural cells, as seen in adults (Flici et al., 2017). Instead, *Soxb2* downregulation caused a loss of RFamide^+^ neurons but not of GLWamide^+^ neurons. Because GLWamide^+^ neurons are essential for metamorphosis of *Hydractinia* larvae (Plickert et al., 2003), it is possible that the animals preferentially generate this type of neuron when progenitors are limited. Conversely, nematoblast numbers increased upon *Soxb2* downregulation, suggesting that Soxb2 normally inhibits nematogenesis in embryos, in contrast to its function in adults (Flici et al., 2017).

Despite being expressed in embryos and larvae (Flici et al., 2017), no visible phenotype was observed following *Soxb3* downregulation, other than the effect of its downregulation in adults (Flici et al., 2017). Therefore, *Soxb3* may not play a distinct role in embryonic neurogenesis, or that the effect of its downregulation in embryos is too subtle to be observed by our methods.

i-cells are concentrated in the lower body column (Bradshaw et al., 2015), neural progenitors are scattered in the mid body column, while most neurons and nematocytes are found in the head. This implies that neural cells differentiate while migrating from the base of the polyp towards the head, reflected by the partly overlapping expression domains of SoxB genes that mark the stages of differentiation. A major question emanating from this work is whether migration is required for neurogenesis cell autonomously, or whether migrating stem cells become exposed to different microenvironments along the oral-aboral axis that drive their differentiation non-cell autonomously.

The poor conservation of SoxB gene sequences across animals make direct functional comparisons difficult. However, the requirement for multiple SoxB genes and migration seem to be common hallmarks of neurogenesis in all studied animals. Additional common features remain to be discovered. *Hydractinia*’s continuous, predictable, and accessible neurogenesis across all life stages provides an excellent model to address these questions.

## ACKNOWLEDGEMENTS

We thank Áine Varley for animal care, and the NIH Intramural Sequencing Center (NISC) for generating the sequence data. Confocal images were taken at the Centre for Microscopy and Imaging Core Facility at NUI Galway. All flow cytometry and imaging cytometry analyses were performed in the Flow Cytometry Core Facility at NUI Galway. Funding: UF is a Wellcome Trust Investigator in Science (grant no. 210722/Z/18/Z, co-funded by the SFI-HRB-Wellcome Biomedical Research Partnership). This work was also funded by a Science Foundation Ireland Investigator Award to UF (grant no. 11/PI/1020), by the NSF EDGE program (grant no. 1923259 to CES and UF), and by the Intramural Research Program of the National Human Genome Research Institute, National Institutes of Health to ADB (ZIA HG000140). SGG was a Marie Curie Incoming International Fellow (project 623748) and was also supported by a Science Foundation Ireland SIRG award (grant no. 13/SIRG/2125). MSS is a Human Frontier Science Program Long-Term Postdoctoral Fellow (grant no. LT000756/2020-L). F was a Hardiman Scholar and also supported by a Thomas Crawford Hayes Research Grant.

## Author contributions

EC and UF initiated and conceptualized the project. UF supervised the project. EC performed laboratory experiments and analyzed data. HF, JMG, and MSS performed experiments. EC and SH performed flow cytometry. EC and KT performed live imaging. MK established nematocyst lectin staining. ETM generated the anti-Piwi2 antibody. SGG performed single cell analysis. PG, CES, and ADB performed the computational analyses. EC and UF wrote the manuscript.

## RESOURCE AVAILABILITY

### Data and code availability

The accession number for the RNA-seq datasets generated in this study is Sequence Read Archive (SRA): BioProjects PRJNA549873 (bulk RNA-seq) and PRJNA777228 (single cell RNA-seq). Accession numbers for each sample are listed in Table S2. The *Hydractinia symbiolongicarpus* genome is available through the NIH National Human Genome Research Institute (https://research.nhgri.nih.gov/hydractinia/). Corresponding data is archived in the NCBI Sequence Read Archive (SRA) under BioProject PRJNA807936.

---

**Figure S1.**
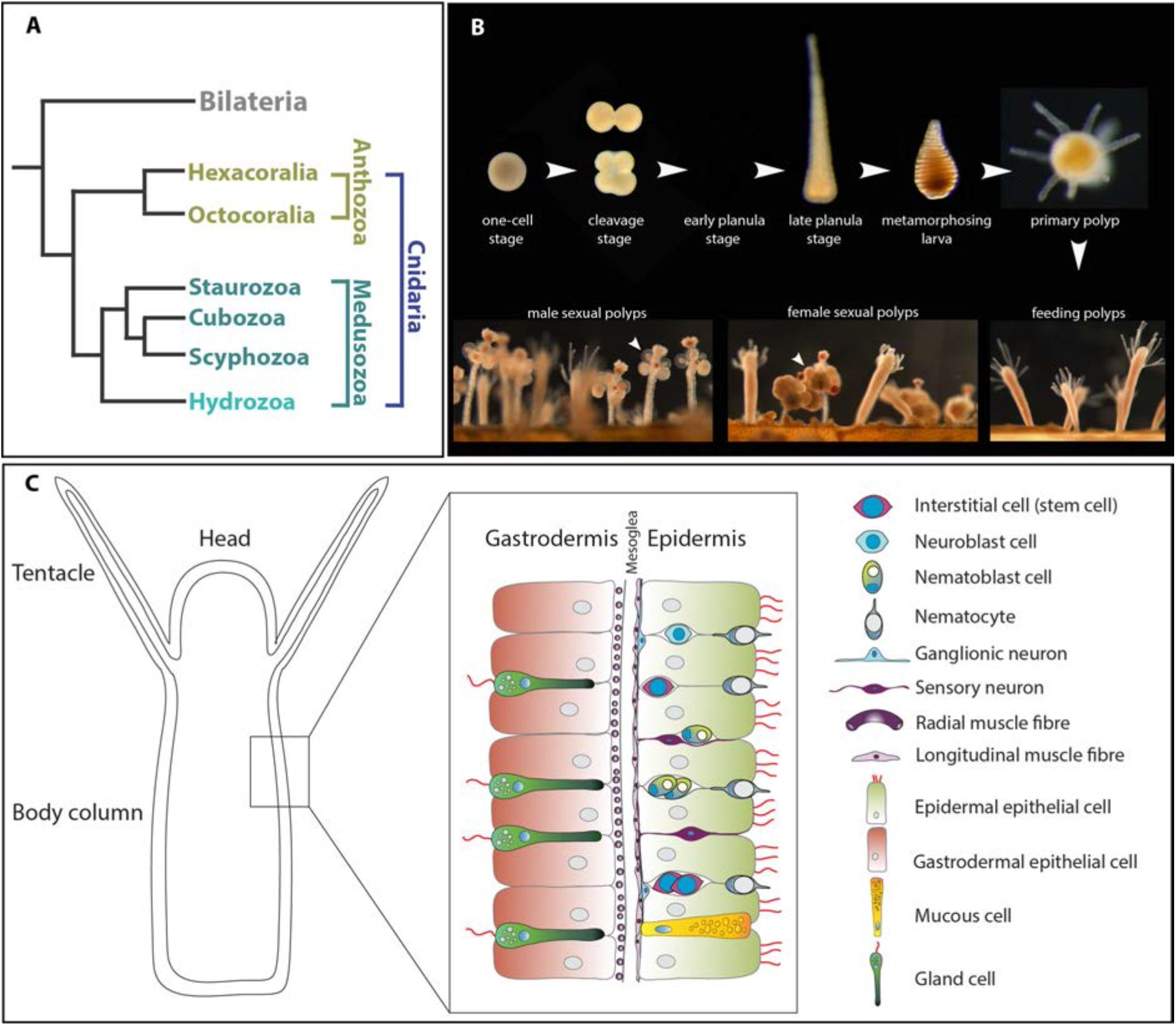
The animal model *Hydractinia*. (A) Simplified cladogram showing the position of various cnidarian clades relative to bilaterians. (B) The life cycle of *Hydractinia*. (B) Schematic representation of *Hydractinia*’s body wall and the cell types composing it.

**Figure S2.**
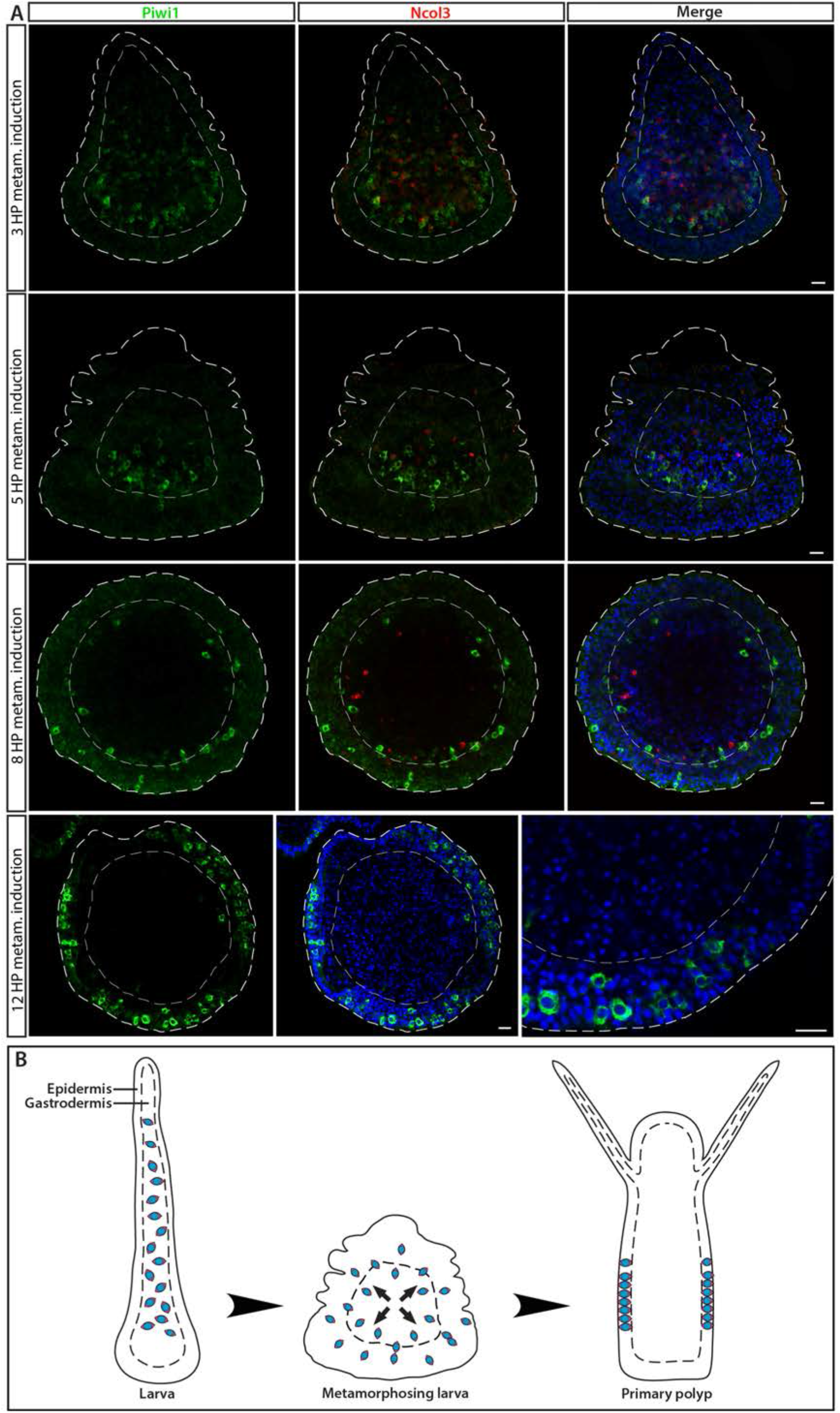
i-cell location during different stages in ontogeny, marked by Piwi1. (A) During embryogenesis and in the larval stage, i-cells are exclusively present in the endoderm; they migrate to the epidermis during metamorphosis. (B) Schematic representation.

**Figure S3.**
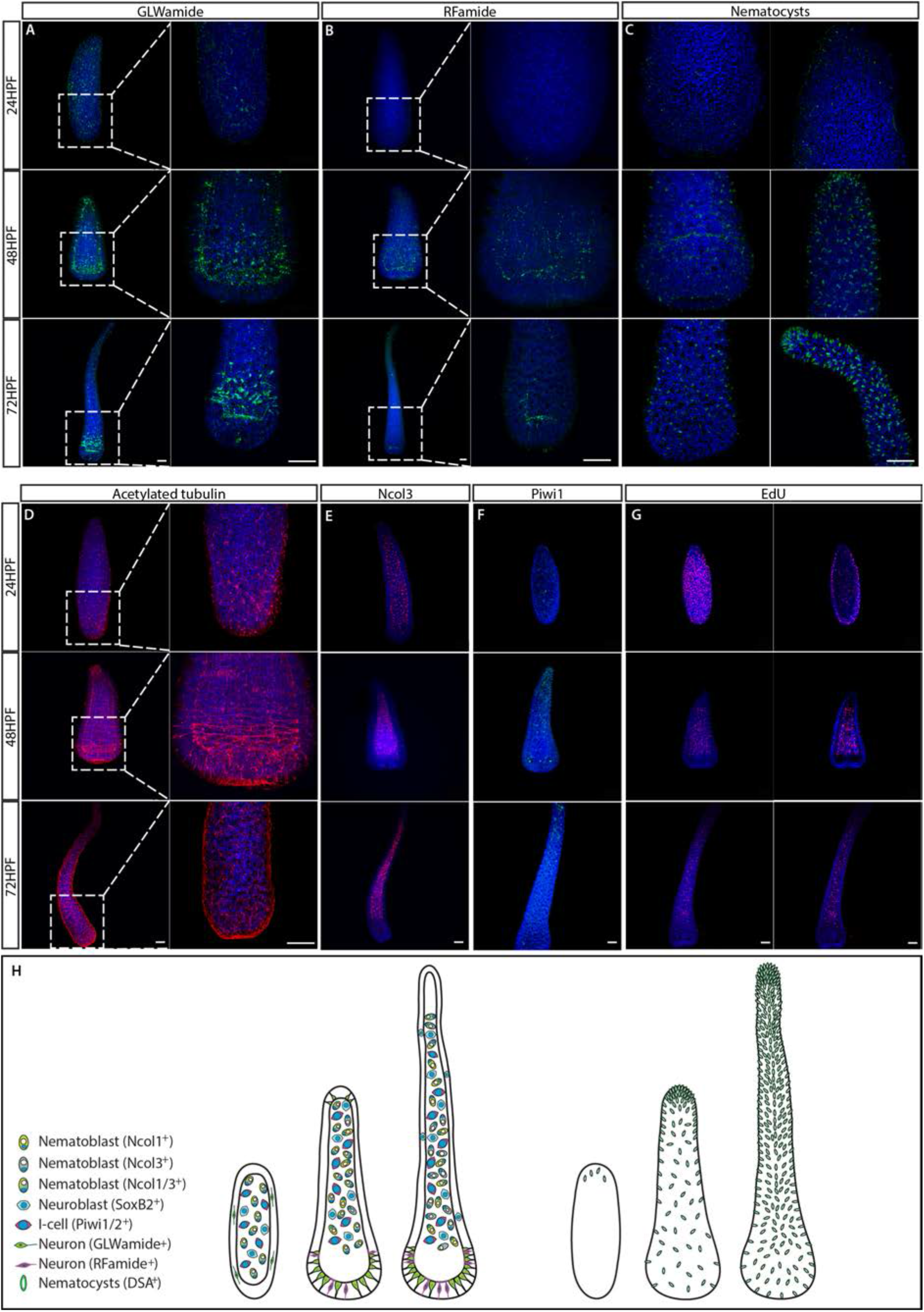
The structure of the nervous system in embryogenesis and larvae. (A) GLWamide^+^ neurons. (B) RFamide^+^ neurons. (C) Nematocyst capsules, representing mature nematocytes, visualized by lectin staining. (D) Acetylated tubulin immunostaining . (E) Ncol3+ nematoblasts. (F) Piwi1+ i-cells. (G) S-phase cells. (H) Schematic representation showing 24 hour embryos, 2 day pre-panula, and 3 day planula larva that is metamorphosis competent.

**Figure S4.**
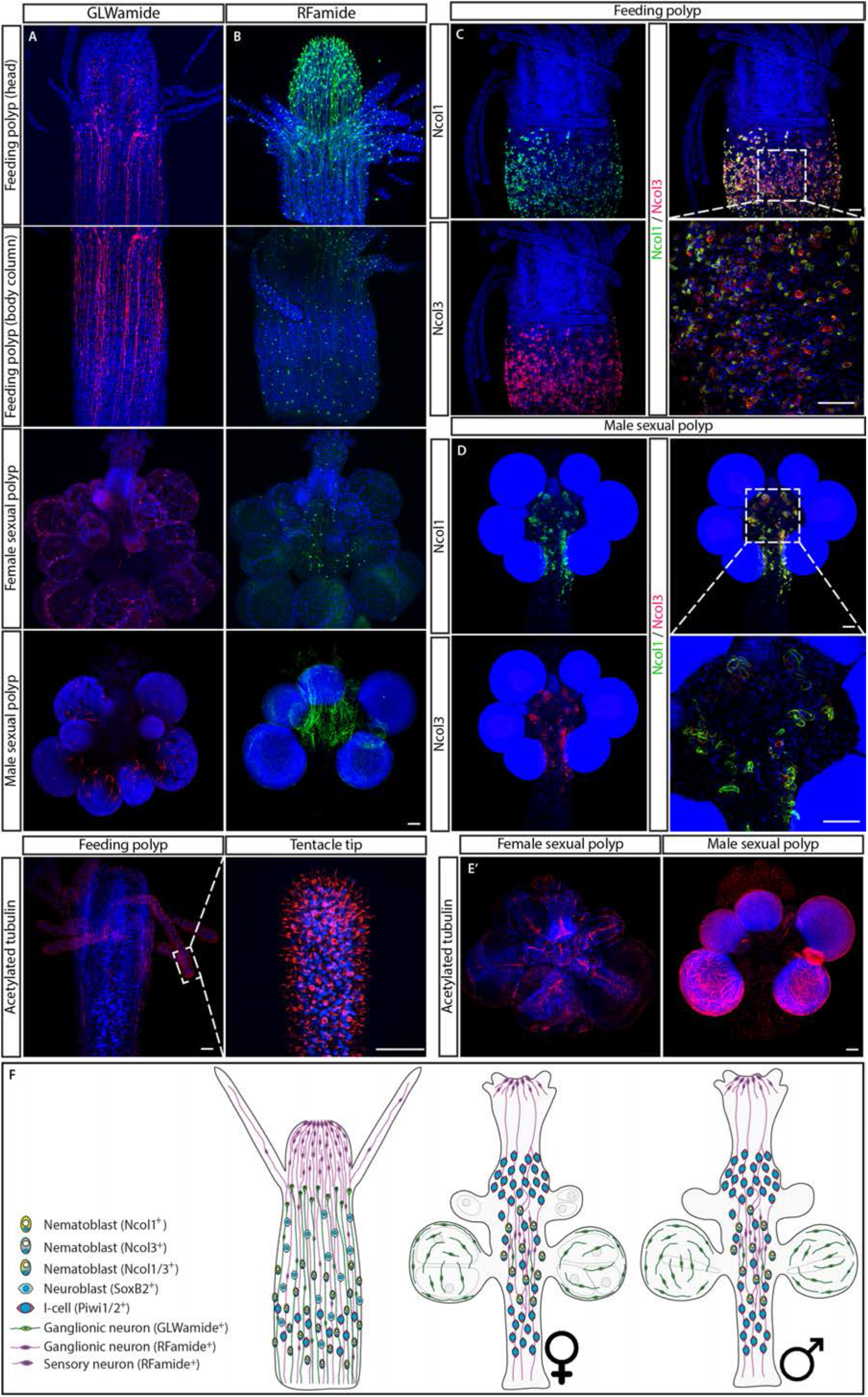
The structure of the nervous system in adult feeding and sexual polyps. (A) GLWamide^+^ neurons. (B) RFamide^+^ neurons. (C) Ncol1+ and Ncol3+ nematoblasts in feeding polyps. (D) Ncol1+ and Ncol3+ nematoblasts in sexual polyps. (E) Acetylated tubulin immunostaining in sexual polyps. (F) Schmatic representation. Scale bars = 40 μm

**Figure S5.**
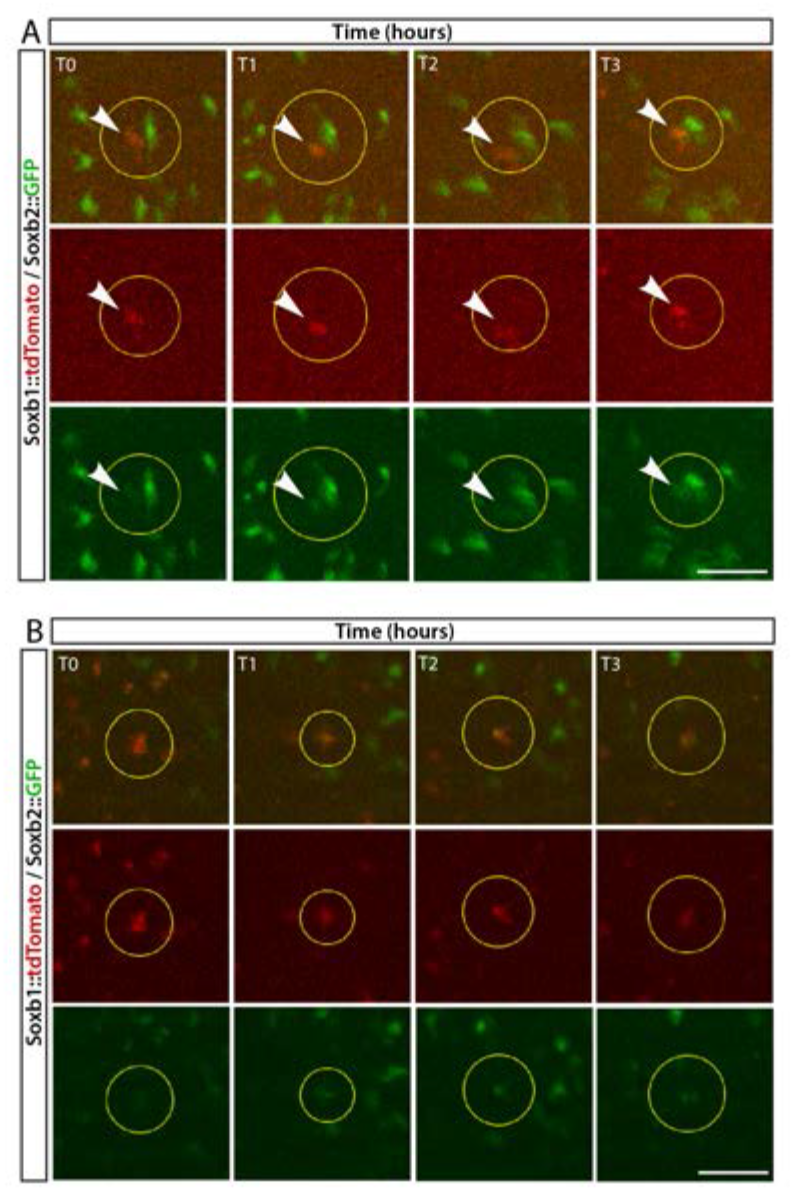
*In vivo* time lapse imaging of a *Soxb1::tdTomato/Soxb2::GFP* double transgenic reporter animal. (A, B) Two cases of gradual upregulation of the *Soxb2* reporter in *Soxb1* expressing cells over an 8 hour time frame.

**Figure S6.**
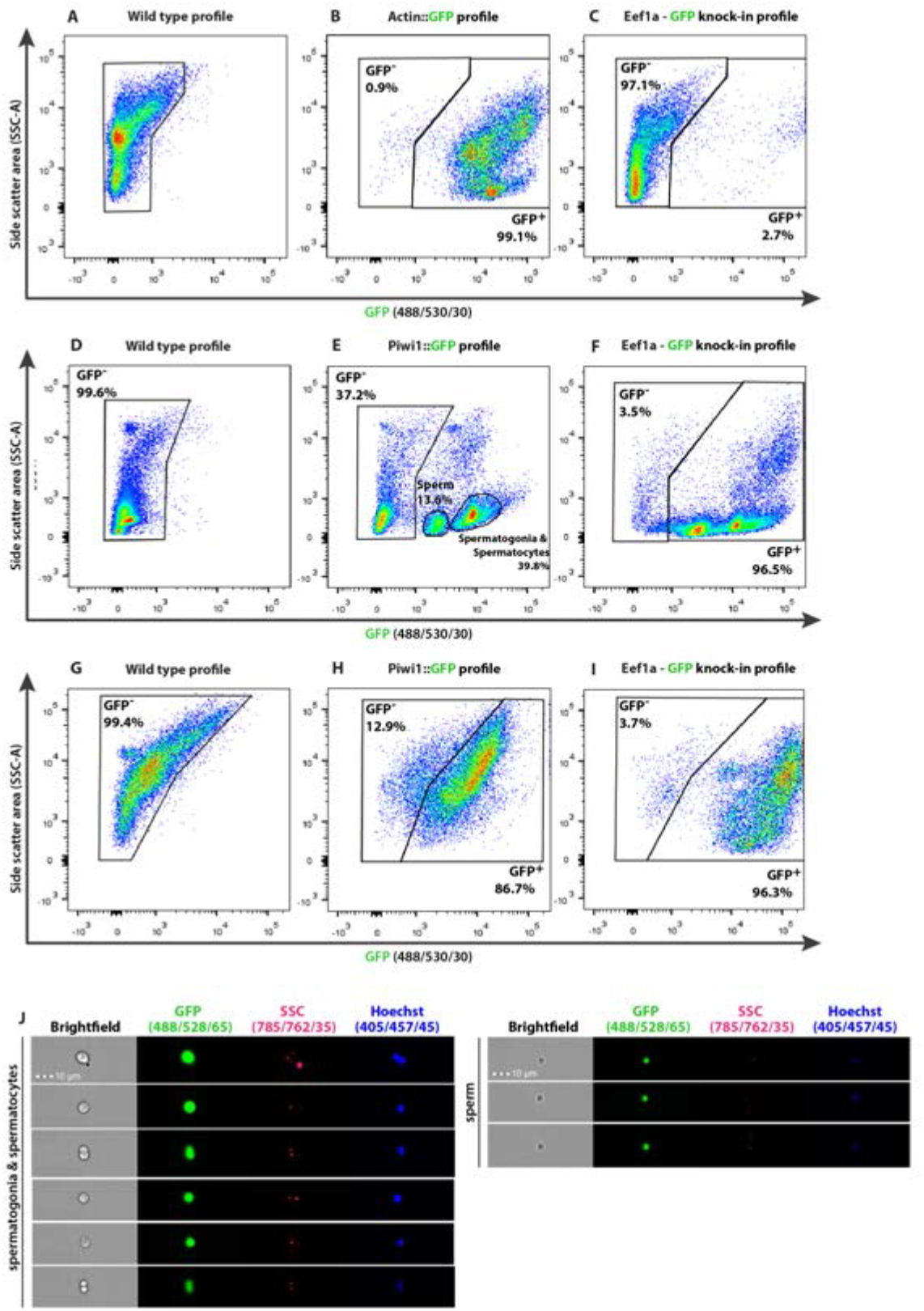
Representative flow cytometry density plots identifying subpopulations as defined by internal complexity (Side Scatter (SSC)) and level of GFP expression of *Hydractinia* transgenic reporter animals and matched wild type animals (A-I) and imaging flow cytometry (J). (A-C) Characterization of feeding polyps, (D-F) male and (G-I) female sexual polyps. The nature of the reporter transgene is indicated above each density plot. (J) Imaging flow cytometric analysis of dissociated male sexual polyps of *Piwi1::GFP* reporter line, displaying the different stages of spermiogenesis including mature sperm cells. Scale bars =10 μm.

**Fig S7.**
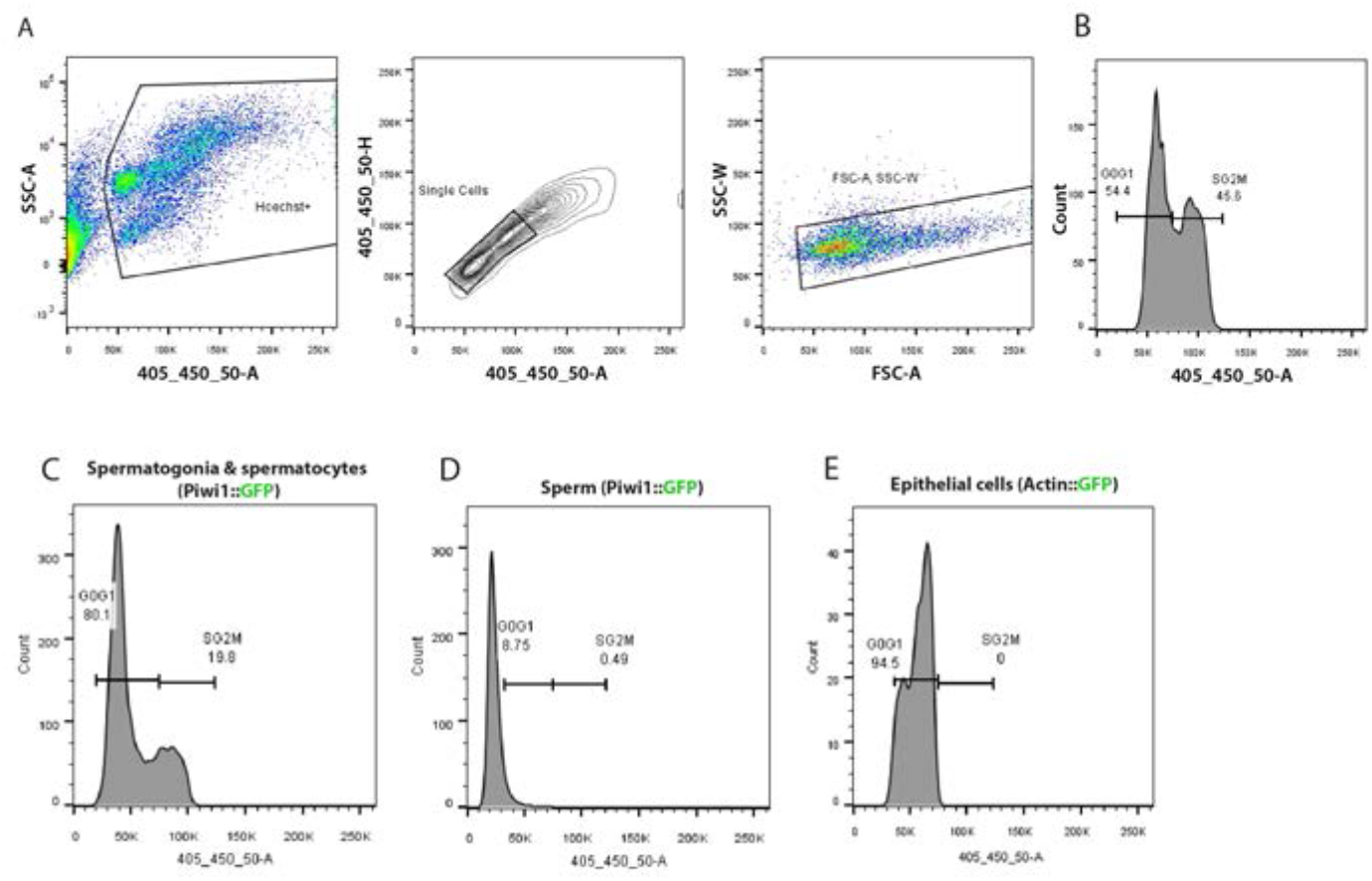
Typical flow cytometry gating strategy used for cell cycle analysis of transgenic animals and a wild type animal. (A) Density plot of cells stained with Hoechst 33342 cell permeant dye versus side scatter (SSC-A). The gated population represents Hoechst positive live, nucleated cells. Doublets are subsequently discriminated by Hoechst Area verus Hoescht Height (contour plot) and Forward scatter area (FSC-A) versus Side scatter width (SSC-W) density plot. (B) Histogram showing a typical cell cycle profile of Hoechst labelled cells disassociated from a wild type *Hydractinia* animal. Cell cycle profile of cell subpopulations identified in the male sexual polyps of Piwi1::GFP reporter line (C) and (D) Cell cycle profile of post meiotic sperm characterized by a haploid genome. (E) Cell cycle profile of Epithelial cells isolated by FACS from an epithelial reporter animal.

**Figure S8.**
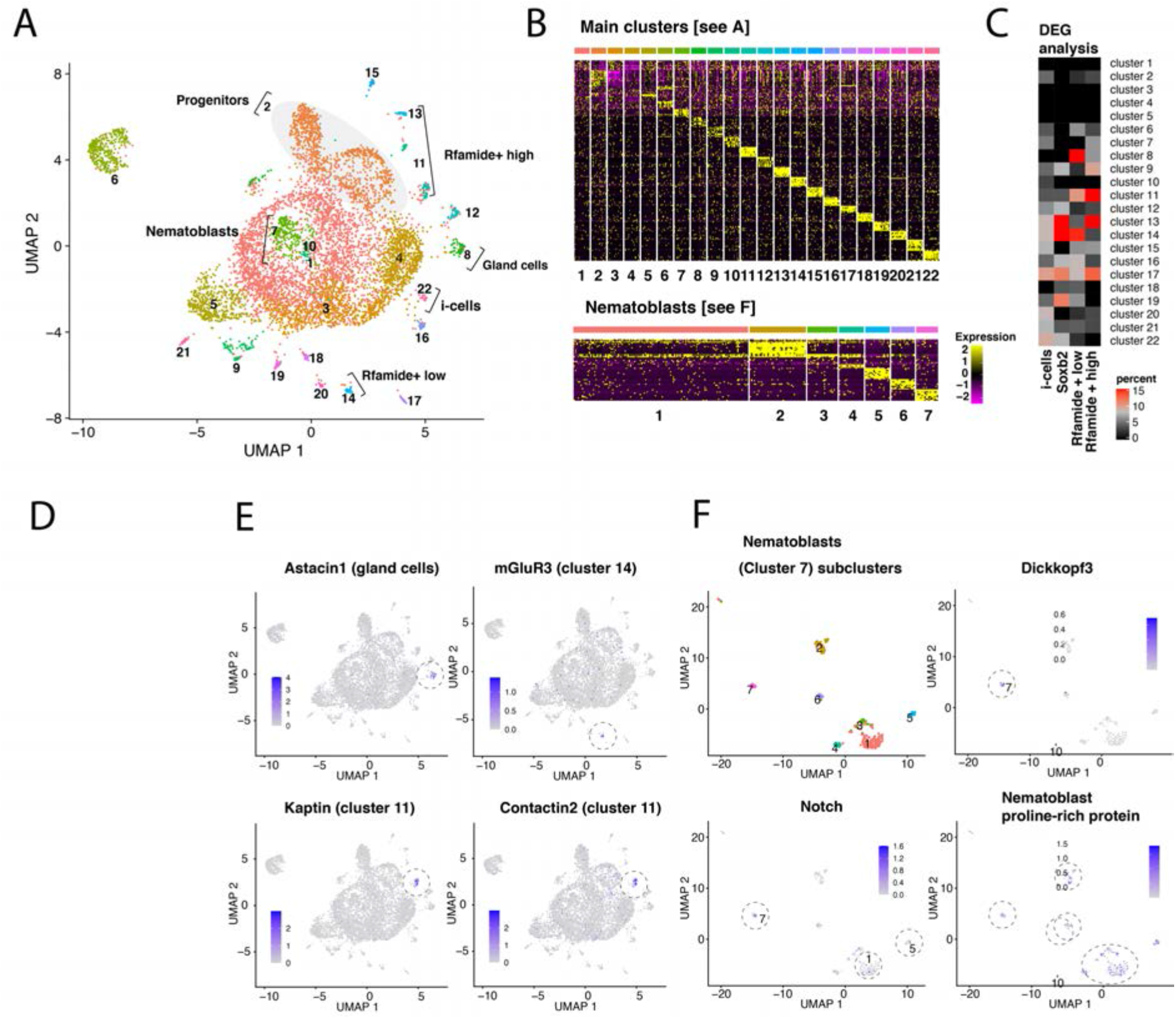
(A) UMAP dimensional reduction projection of 7,071 *H. symbiolongicarpus* cells. (B) Heatmap showing the top ten markers for each cell cluster across all clusters and within nematoblast (Cluster 7) subclusters following Seurat analysis. (C) Heatmap showing the percentage overlap between single-cell RNA cluster markers and differentially upregulated genes in FACS-sorted cells from transgenic reporter animals. The heatmap shows that the overlap between *Soxb2*^+^ sorted cells and clusters 13 and 14 is highest, i.e consistent with *Soxb2* being most highly expressed in these clusters. (D) Dotplot depicting both the percentage proportion and average log-normalised expression of *Soxb1*, *Soxb2* and *Soxb3* genes in all cells across the main clusters as shown in [A]. We find that both soxB1 and soxB2 are expressed in a subset of i-cells; presumably in transition to become soxB2 positive progenitor cells (clusters 13 and 14 in [A]), while soxB3 is only expressed in few cells in cluster 18 (see [A] and [D]). It also appears that subpopulations of neural cells in *H. symbiolongicarpus* transition from a soxB2/RFamide low state (cluster 14) that express a G-protein coupled receptor for glutamate [see E] to a soxB2/RFamide high state (cluster 13), of which some eventually become soxB2 negative while still expressing high levels of RFamide (cluster 11); presumably as terminally differentiated axonal neurons as determined by the markers Kaptin and Contactin2, which are important in synapse and axon formation, maintenance and function (see [E]). This is consisten with our model and bolsters the role of soxB2 as a marker for a neural progenitor cells. (E) UMAP dimensional reduction projections of relevant marker gene expression in representative cluster as annotated in panel (A). (F) Subclustering of nematoblasts (Cluster 7 in panel A): UMAP dimensional reduction projection of 259 cells of *H. symbiolongicarpus* nematoblasts accompanied by projections showing the expression of nematogenesis marker genes (*Dickkopf3*, *Notch* and a proline-rich protein homologous to the nematoblast-specific protein nb039a-sv15 of *Hydra vulgaris*) across several subclusters.

**Figure S9.**
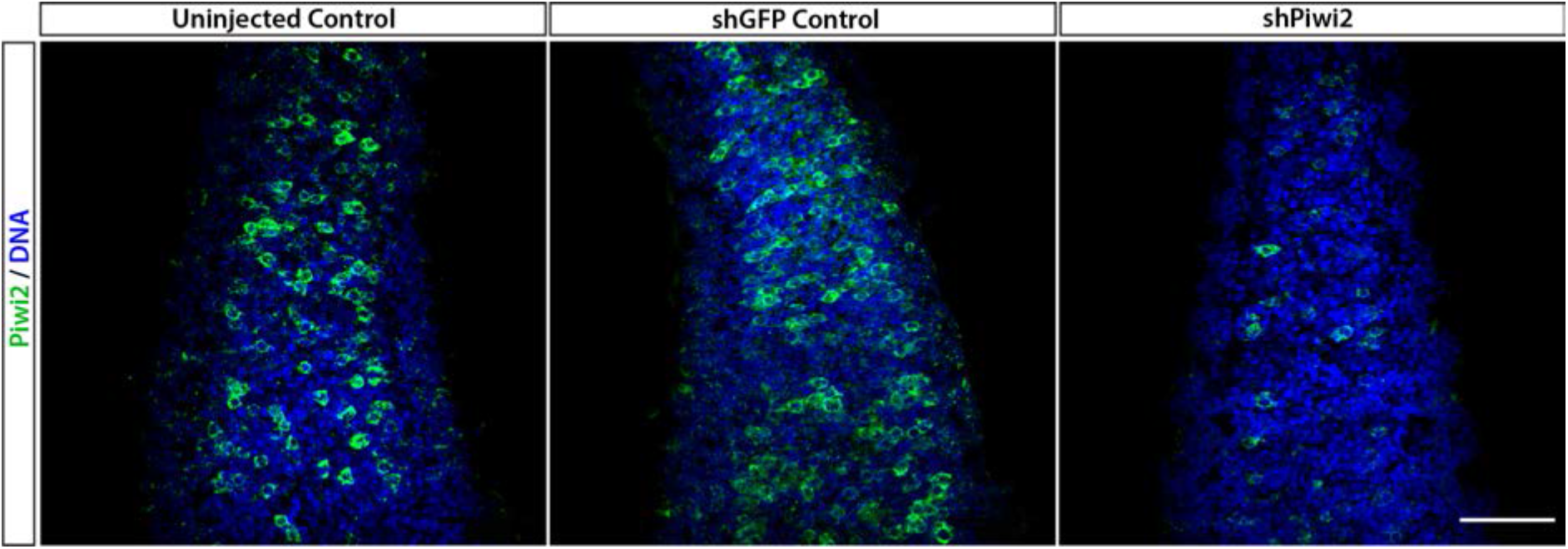
Piwi2 antibody validation. Zygotes were injected with a shRNA targeting *Piwi2* (shPiwi2) or a control shRNA targeting the GFP sequence (shGFP). At 48 hours post fertilization, animals were fixed and stained with the anti-Piwi2 antibody. Fewer Piwi2^+^ i-cells are detectable in shPiwi2 than in the control.

**Figure S10.**
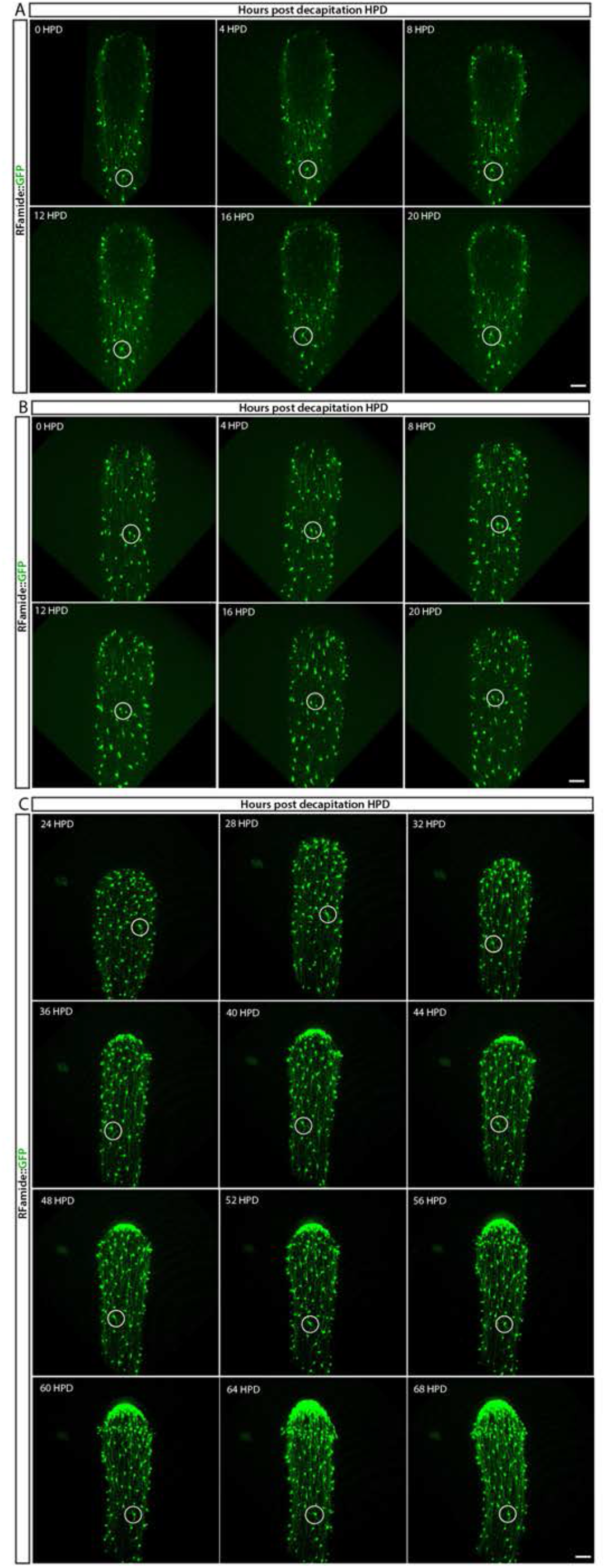
*In vivo* tracing of RFamide-GFP+ neurons during regeneration located in the (A) lower part and (B) upper part of the body column during the first 20 hours post decapitation (HPD), and (C) from 24 HPD until 72HPD. No cell proliferation or migration of individual neurons (circled) was observed, indicating no role during this process. Images are single optical slices. Scale bars = 40μm.

**Figure S11.**
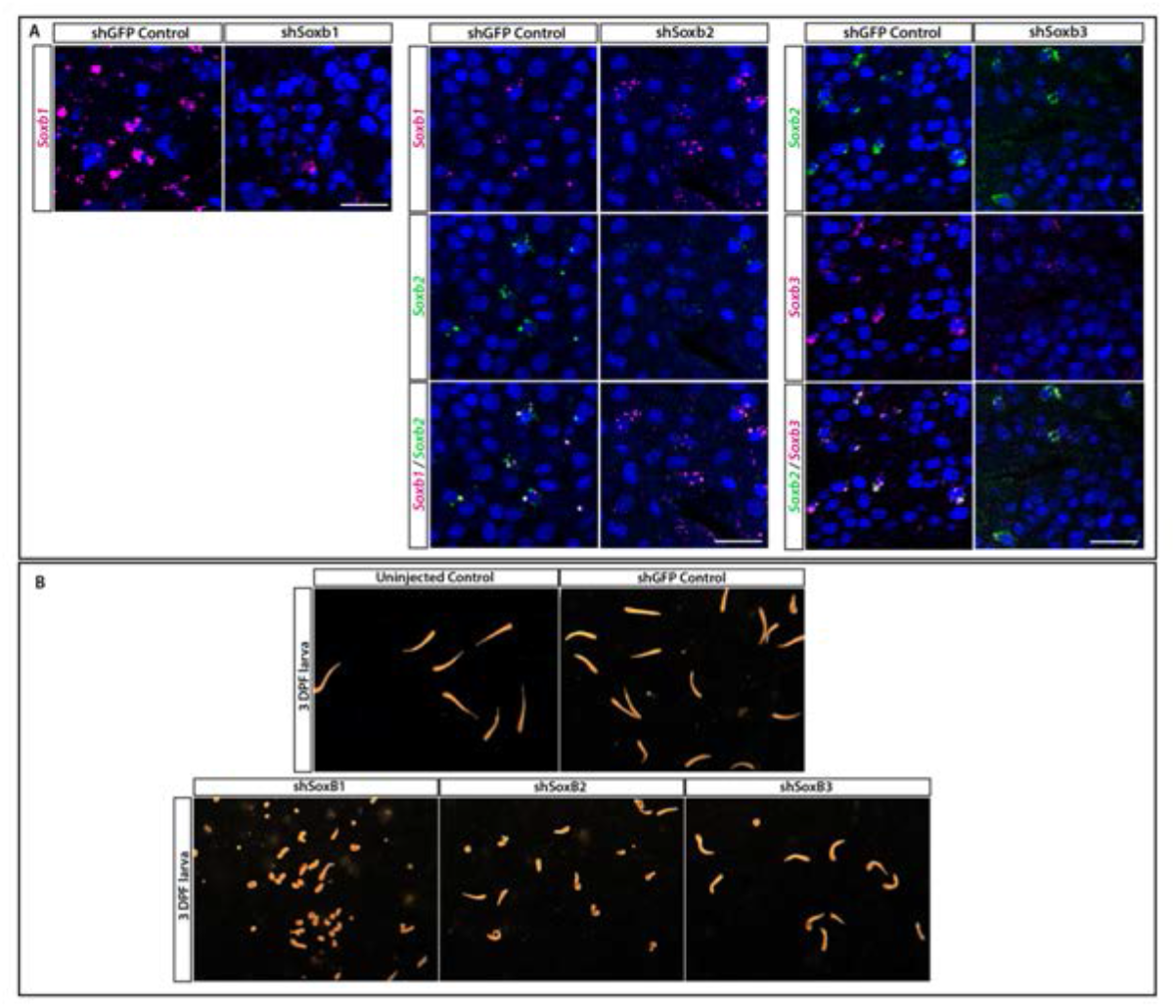
(A) Validation of shSoxB1, shSoxB2 and shSoxB3 knockdowns by single molecule fluorescence in-situ hybridization (SABER FISH) on 3DPF larvae. Larvae injected with shSoxB1 showed lower expression levels of SoxB1 compared to the control animals. Larvae injected with shSoxB2 showed lower expression levels of SoxB2 compared to the control animals and SoxB1 levels remained unaffected. Larvae injected with shSoxB3 showed lower expression levels of SoxB3 compared to the control animals and SoxB2 levels remained unaffected. Images are single optical slices. Scale bars = 40 μm. (B) Morphology of control, shGFP-injected, shSoxB1-injected, shSoxB2-injected, and SoxB3-injected larvae. Scale bar: 498μm. (C) Forced expression of *SoxB1::GFP* in a neuronal context was lethal. Remnants of GFP could be seen in the vacuoles of phagocytic cells. (1) Polyp head and body. (2) Part of stolonal tissue. GFP was identified by direct fluorescence; Ncol1 by IF. Images are z-stack maximum projections. Scale bars = 40 μm.

**Figure S12.**
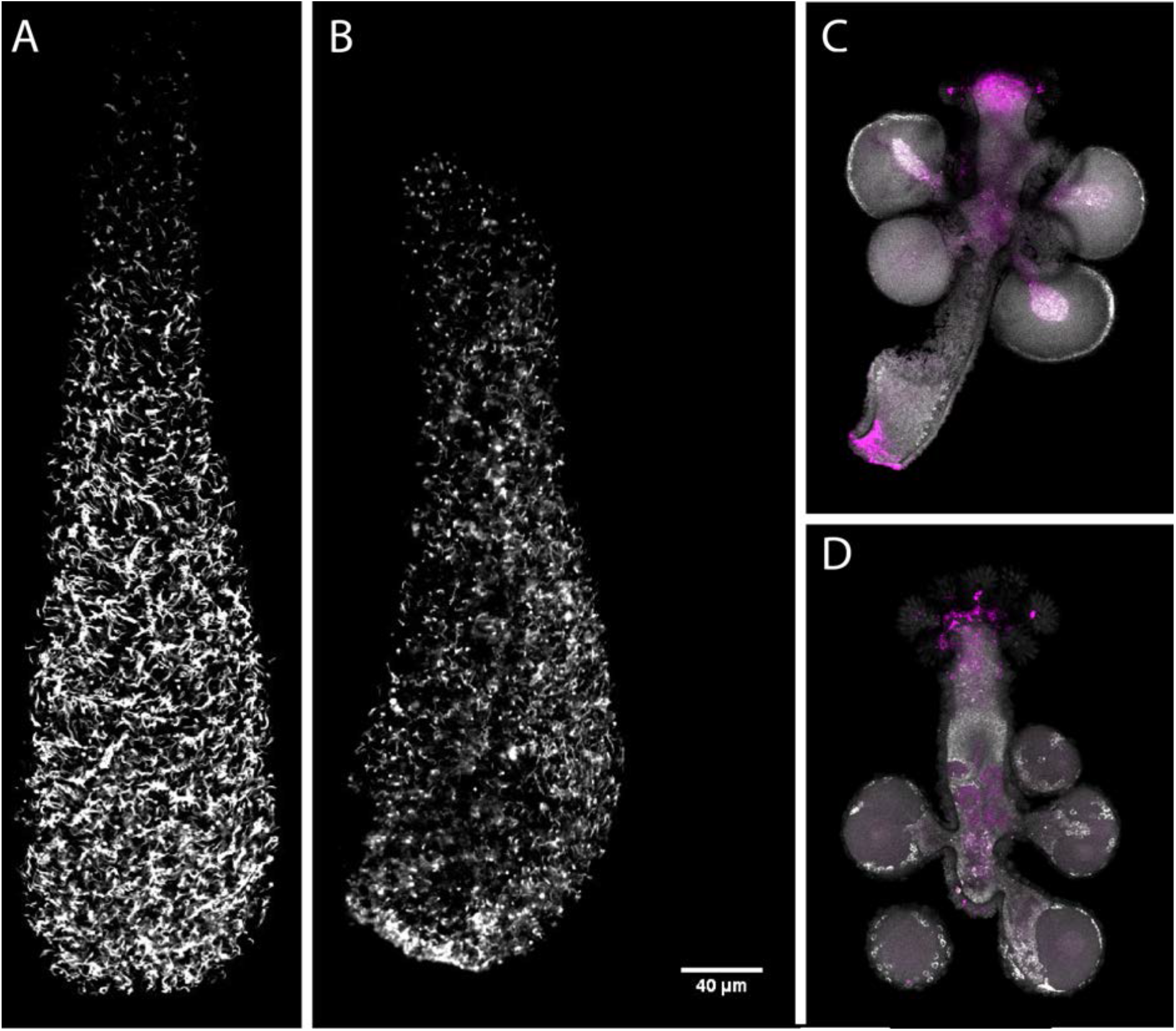
Effect of *Soxb1* downregulation on epidermal ciliation in 3-day old larvae using acetylated tubulin antibody staining and in situ hybridization of *Soxb2* in sexual polyps (A) shGFP-injected animal. (B) shSoxb1-injected animal. (C) Male sexual polyp. (D) Female sexual polyp.

